# Non-canonical ORFs are an important source of tumor-specific antigens in a liver cancer meta-cohort

**DOI:** 10.1101/2023.10.30.564375

**Authors:** Marta E. Camarena, Patrick Theunissen, Marta Ruiz, Jorge Ruiz-Orera, Beatriz Calvo-Serra, Robert Castelo, Pablo Sarobe, Puri Fortes, Júlia Perera-Bel, M. Mar Albà

## Abstract

The expression of tumor-specific antigens during cancer progression can trigger an immune response against the tumor. Antigens that have been used as cancer vaccines are those originated by non-synonymous mutations and those derived from cancer/testis antigens. However, the first class is predominantly patient-specific, preventing the development of therapies than can benefit multiple patients, and the second one offers a limited set of actionable targets. A possible alternative is the use of peptides derived from non-canonical ORFs (ncORFs). While many ncORFs have been shown to be translated in cancer cells, their tumor-specificity and patient distribution remains to be determined. Here we analyze RNA sequencing data 117 hepatocellular carcinoma (HCC) tumors and matched healthy tissue, together with ribosome profiling data from an additional 10 HCC tumors, to answer these open questions. Combining HLA-epitope binding predictions and experimental validation experiments we conclude that around 40% of the tumor-specific antigens in HCC are likely to be derived from ncORFs in lncRNAs, including two peptides that can trigger an immune response in mice. We identify a subset of 33 tumor-specific lncRNAs expressing novel cancer antigens shared by more than 10% of the HCC analyzed, which could be combined to target a large proportion of the patients. The results of the study open new avenues for extending the range of anti-cancer vaccines.

## INTRODUCTION

Immunotherapy approaches against cancer, including immune checkpoint inhibitors (ICI) and vaccines, rely on the ability of the immune system to recognize ‘non-self’ antigens bound to HLA receptors. Such neoepitopes can originate from non-synonymous mutations in the cancer genome that result in mutated peptides, but also from aberrant gene expression in tumors. The first class of antigens is especially relevant in cancers associated with a large number of mutations, such as melanoma, lung or bladder cancer ^1^. Not surprisingly, tumor mutational burden and the number of mutated peptides with predicted affinity to HLA receptors are positively correlated with the response to ICI ^2–4^.

The second class of antigens might be particularly relevant to develop therapeutic strategies for tumors that mutate less frequently, such as hepatocellular carcinoma (HCC), which represents ∼90% of cases of liver cancer. Known cancer-specific antigens include the so called cancer/testis antigens (CTA), as well as peptides derived from reactivated human endogenous retroviruses (HERVs) ^5,6^. These antigens can be found in different cancer types and they can be shared by several patients. Some of them, such as MAGE1A and NY-ESO, have been the basis of several cancer vaccines ^7^. Current limitations are the relatively low number of suitable targets with high tumor specificity and their sparse expression in cancer patient samples.

A promising approach to expand the current range of cancer-specific antigens that can be targeted by immunotherapy approaches is to consider the translation products of non-canonical ORFs (ncORFs). These ORFs are located in sequences that are not annotated as protein-coding. One well-studied example are the MELOE-1 and MELOE-2 peptides encoded by the long non-coding transcript *meloe* ^8,9^. MELOE-1 generates a CD8+ T cell epitope, and the adjuvant injection of lymphocytes containing MELOE-1 specific CD8+ T cells to melanoma patients has been shown to be beneficial ^10^. In the past few years, thousands of lncRNAs containing translated ncORFs have been described ^11–15^. Additionally, mass spec immunopeptidomics data from cancer cell lines and tumors indicates that ncORFs can generate peptides that are presented by HLA receptors ^16–19^. It has been reported that ncORF products can represent up to 15% of the HLA-I bound peptides in certain tumor types ^20^, a sizable fraction that remains largely uncharacterized. In addition, they appear to give rise to HLA-I bound peptides more frequently than standard proteins ^21^.

To be able to avoid immune self-tolerance, the ncORFs need to be expressed in a tumor-specific manner. But, due to the lack of studies comparing tumor and healthy tissues from the same set of patients, it is unclear how many of the previously reported ncORF-derived antigens are actually restricted to tumors. Thus, it is not known if peptides derived from ncORFs could be relevant as therapeutic targets. To address these questions, we have focused on tumor and matched healthy tissue sequencing data from a larger number of hepatocellular carcinoma (HCC) patients. Treatment of HCC in advanced stages remains a challenge ^22^. Because this is a type of cancer with relatively few mutations, antigens derived from tumor-specific transcripts could play a major role in driving immunogenicity. We present new data supporting that ncORFs are a relevant source of tumor-specific antigens in HCC. The findings could have important implications for the development of cancer vaccines of wide applicability.

## RESULTS

### Analysis of tumor and normal matched transcriptomics data

We identified four HCC patient cohorts with transcriptomics data for both tumor and adjacent normal tissue (Figure 1a HCC1-HCC3 and TCGA, Table S1) ^23–26^. Altogether, this represented a meta-cohort of 117 patients. We also identified ribosome profiling (Ribo-seq) sequencing data from an additional set of 10 HCC tumors (HCC4)^26^. We used several previously described HCC biomarkers to validate these datasets: two genes that tend to be over-expressed in HCC – TERT (Cancer Genome Atlas Research Network 2017) and THBS4 ^27^ - and one that is usually under-expressed - MT1M ^28^. Consistent with previous findings, we found that TERT and THBS4 had significantly higher expression levels in tumor than in normal matched samples in all cohorts, and that MT1M showed the opposite tendency (Figure 1b).

**Figure 1.**
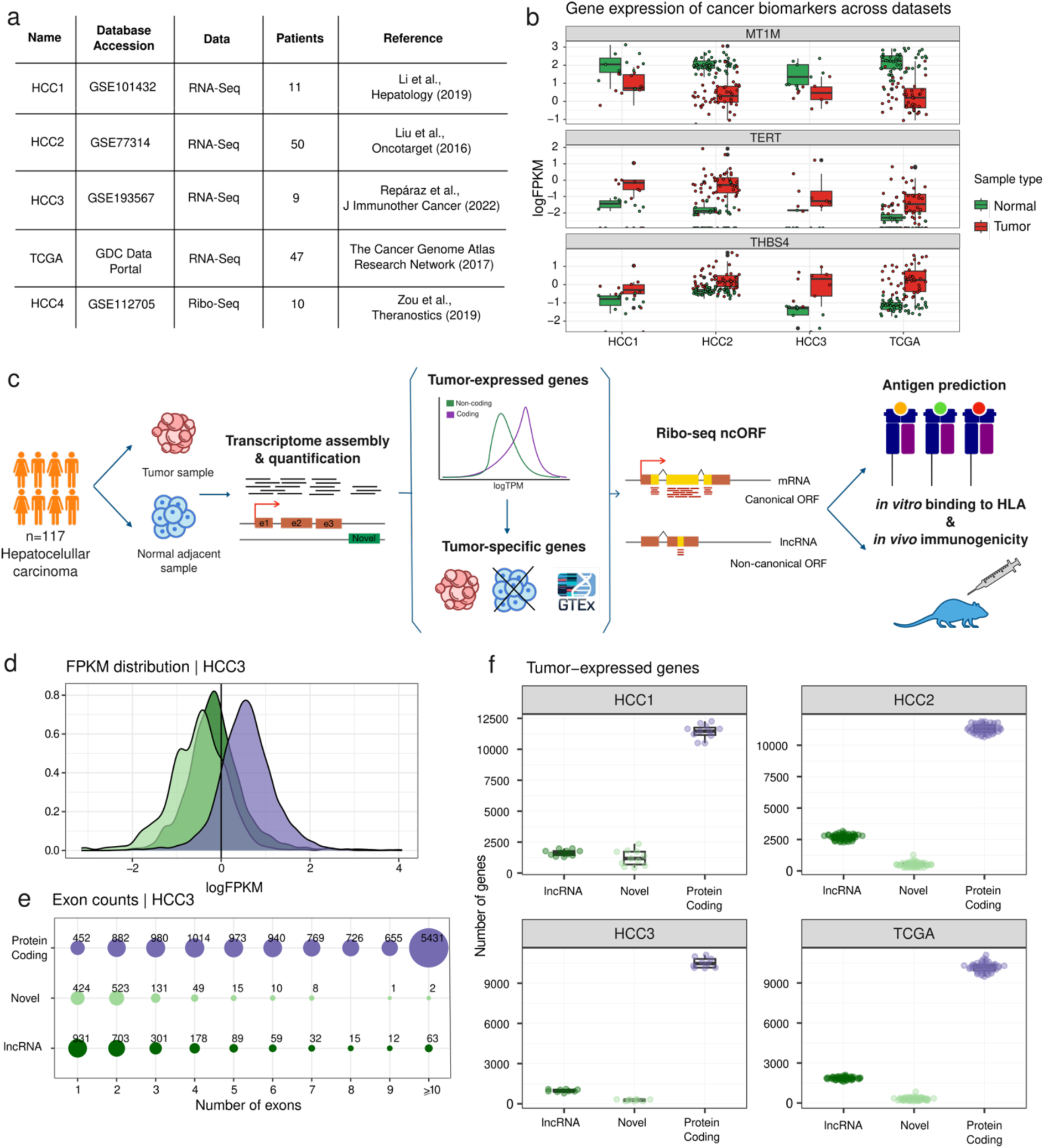
The hepatocellular carcinoma (HCC) transcriptome. a. Datasets used in the study. A total of 117 HCC patients with matched tumor-adjacent tissue RNA-Seq data were analyzed (datasets HCC1-HCC3 and TCGA). An additional cohort comprising 10 patients with matched tumor-adjacent tissue Ribo-Seq data (HCC4) was used to measure translation of ncORFs in HCC. **b. Biomarkers of HCC in the four datasets**. Gene expression was measured as fragments per kilobase per million reads (FPKM), in both tumor and adjacent tissue samples (normal). By paired Wilcoxon signed-rank test, we confirmed that the expression of MT1M, TERT and THBS4 coding genes was significantly different in tumor samples and in adjacent tissue following the expected trends (MT1M p-value = 1.596e-19; TERT p-value = 1.578e-19; THBS4 p-value = 9.941e-20). **c. Main steps of the computational and experimental pipeline.** We gathered RNA-Seq and Ribo-Seq data from matched tumor/normal samples. We quantified gene expression and reconstructed non-annotated transcripts. We then determined the tumor-specific transcriptome in each patient. We predicted the translation rate of lncRNAs and novel transcripts. We quantified the generation of tumor-specific antigens derived from ncORFs versus other sources and performed experiments to validate HLA-binding and immunogenicity. **d. Distribution of gene expression levels for different types of transcripts**. LncRNAs and novel transcripts were typically expressed at lower levels than protein-coding genes, although there was a considerable overlap in expression levels between the classes. The line at 1 FPKM indicates the expression cutoff used to consider a transcript as expressed. Data shown is for dataset HCC3. **e. Number of exons in different types of transcripts.** As expected, lncRNAs and novel transcripts tended to have a lower number of exons than coding genes; the data shown is for the HCC3 dataset. **f. Relative abundance of different types of transcripts in tumors.** Coding genes were the largest class of expressed transcripts, followed by lncRNAs and novel transcripts.

After validating the datasets with the above biomarkers, we designed a pipeline that combined different computational and experimental methods to unravel the impact of ncORFs in the generation of tumor-specific antigens in the set of 117 patients (Figure 1c). The first steps were centered on the quantification of gene expression, the discovery of novel transcripts and the identification of tumor- specific transcripts from tumor/normal matched information. We also predicted ncORF translation by the analysis of Ribo-seq data, and putative MHC-I binding peptides using patient-specific HLA information. To validate the predictions, we performed *in vitro* HLA-peptide binding assays of a subset of the candidates, as well as immunogenicity experiments in mice expressing the human HLA receptor (Figure 1c). The analysis provided new information about the quantitative relevance of different types of tumor antigens in HCC. It was also informative on the distribution of these antigens in the patient population. The results of these analyses are described in the next sections.

### Characterization of the full transcriptome of 117 HCC tumor samples

We used the RNA-Seq data from the 4 HCC cohorts to quantify the expression level of protein-coding genes and lncRNAs, as well as to perform genome-guided *de novo* transcript assembly and identify transcripts not annotated in Ensembl ^29,30^. LncRNAs and novel transcripts showed overall lower expression values than protein-coding genes (Figure 1d); only those expressed at significant levels were selected for further analyses (Figure S1, Tables S2-S4). As expected, lncRNAs and novel transcripts tended to have a lower number of introns than protein-coding genes (Figure 1e, Figure S2). We also noted that the majority of novel transcripts, even if not annotated in Ensembl, matched entries in miTranscriptome, a gene database that contains an extended set of cancer transcripts (Iyer et al., 2015)(Figure S3). Each tumor sample expressed around 10,000-12,000 protein-coding genes together with 2,000-4,000 lncRNAs and novel, non-annotated, transcripts (Figure 1f, Figure S4). We found that, in general, the expression of lncRNAs and novel transcripts was more patient-specific than the expression of protein-coding genes (Figure S5).

### Tumor lncRNAs are pervasively translated

Recent studies have shown that many lncRNAs contain ORFs that are translated into small proteins ^11,13–15,31–33^. Here we used Ribo-Seq data from hepatocellular carcinoma (cohort HCC4, Figure 1A) to predict the level of translation of the previously identified tumor lncRNAs and novel transcripts (cohorts HCC1-3 and TCGA). To obtain reliable estimates, we focused on those transcripts that were widely expressed in HCC4 and at least another cohort (see Methods). In addition to ATG, we also considered near cognate codons (ACG, CTG, GTG and TTG) as putative start sites, as these codons have been shown to frequently initiate translation of ncORFs ^18,21,34^. Translation was predicted using RibORF (v 1.0)^13^(Figure 2a)(Figure S6 and Methods). We identified 251 unique lncRNAs translated, including 124 transcripts that were common to all cohorts (Figure 2b). A large fraction of the latter transcripts (86 out of 124) had also been predicted to be translated in a study that analyzed different cancer cell lines and tumors ^18^(Table S5), which reinforced our results. Because the latter study did not include HCC data, this also implies that many of these lncRNAs are expressed in different cancer types.

**Figure 2.**
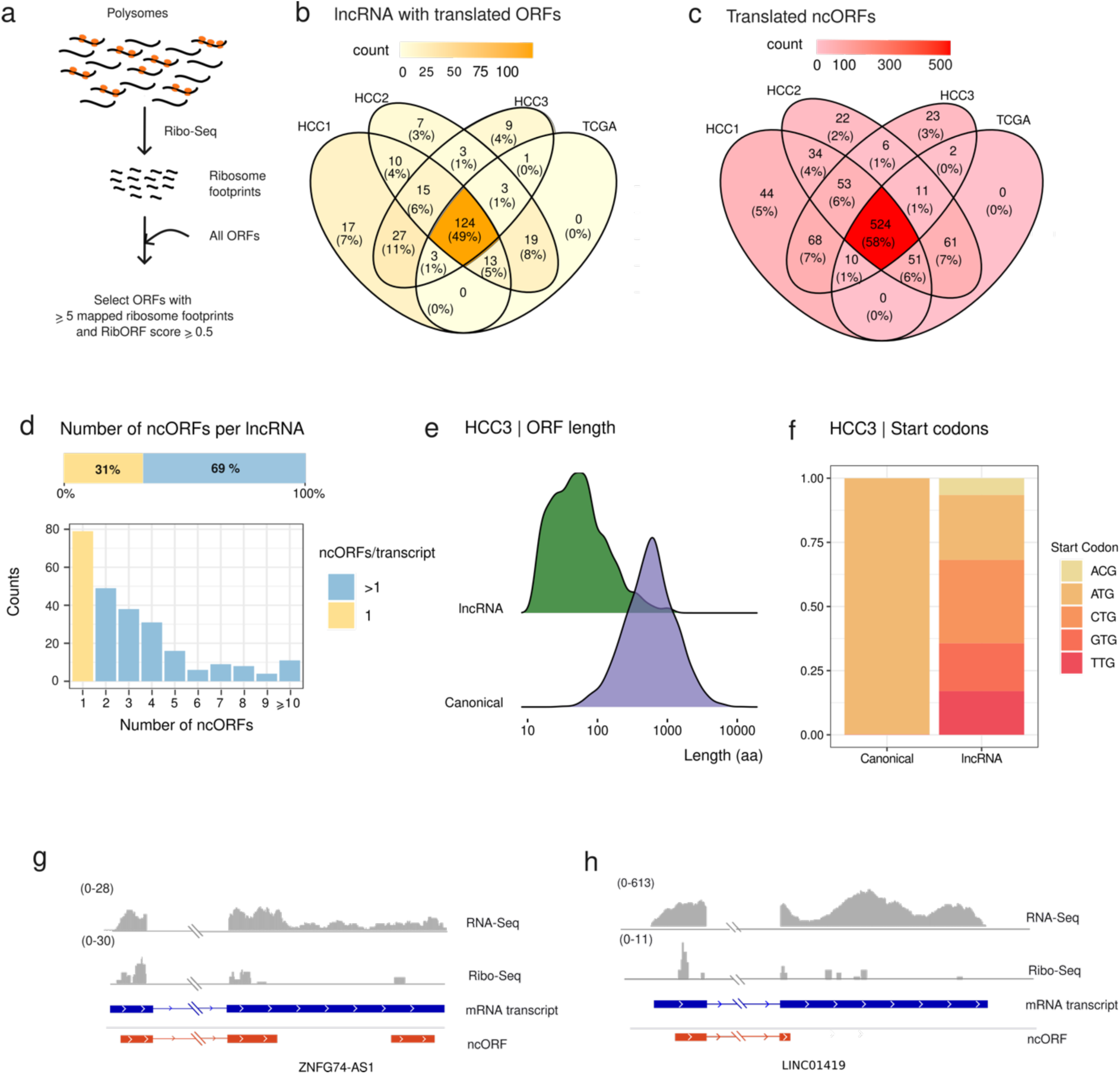
Translation of ncORFs in tumor expressed lncRNAs. a. Prediction of translated ORFS using Ribo-Seq data. From the total predicted non-canonical ORFs, we analyzed translation patterns in ncORFs with at least 5 mapped Ribo-seq reads, selecting those that had a RibORF score of at least 0.5. **b. Comparison of lncRNAs containing ncORFs with signatures of translation from different cohorts**. The intersection between the sets of translated lncRNAs in the four different transcriptomics cohorts is shown. 124 out of a total of 251 translated lncRNAs were shared across all cohorts. **c. Comparison of ncORFs with signatures of translation from different cohorts**. The intersection between the sets of translated ncORFs shown in the different cohorts is shown. 524 out of a total of 909 translated ncORFs were shared across all cohorts **d. Some lncRNAs contain several putatively translated ncORFs.** The graph shows the distribution of the number of translated ncORFs per transcript. From a total of 251 lncRNAs, 79 translate one single ncORF and 172 translate more than one ncORF. **e. ncORFS are significantly smaller than canonical coding sequences.** Comparison of the ORF length distribution of ncORFs versus canonical ORFs, with median values of 39 and 456 aa, respectively. Differences are significant at p-value < 2.2e-16 (Kolmogorov-Smirnoff test). **f. Frequency of different start codons in canonical coding sequences and ncORFs.** ATG as well as ACG, CTG, TTG and GTG were considered as putative start codons. **g. Translation of ZNF674-AS1.** Coverage of RNA-Seq and Ribo-Seq reads, and putatively translated ORFs are indicated. The second exon of the mRNA transcript is shortened for visualization proposes. No Ribo-Seq signal was detected in the region not shown. **h. Translation of LINC01419.** Coverage of RNA-Seq and Ribo-Seq reads, and putatively translated ORFs, are indicated.

The number of ncORFs for which translation was detected was 909, with 524 being common to all cohorts (Figure 2c). Indeed, the majority of the transcripts contained multiple translated ncORFs (Figure 2d). As expected, the resulting proteins tended to be smaller than canonical proteins (Figure 2e). Translation predictions comprised ORFs initiated at ATG but also at alternative sites, especially CTG (Figure 2f).

We used the ribosome profiling data to compute a translation index for lncRNAs and novel transcripts, which we defined as the fraction of ncORF sequence predicted to be translated. In the case of lncRNAs the translation index was 0.116. It was calculated taking into account the total percentage of translated ORFs (8.3%, Table S6) as well as the fact that translated ORFs tended to be somewhat longer than non-translated ORFs (Figure S7). The same estimation for novel transcripts resulted in a much smaller translation index (0.0053), indicating that the latter transcripts are rarely translated.

Figure 2g and 2h show examples of putatively translated ORFs in ZNF674-AS1 and LINC01419, respectively. ZNF674-AS1 is transcribed in antisense direction to the protein-coding gene ZNF674 through the use of a bidirectional promoter and low expression in tumors is associated with bad prognosis ^35^. LINC01419 is a lncRNA of unknown function that is transcribed and translated in a subset of tumor samples but in none of the healthy controls.

### Tumor-specific transcripts are enriched in lncRNAs and novel transcripts

Peptides generated from ncORFs in tumor-specific lncRNAs are a potential source of cancer antigens with immunotherapy applications, as described for some of the canonical cancer/testis antigens ^36^. To determine how many of the transcripts expressed in tumors were tumor-specific, we checked if they were expressed in matched healthy liver samples, GTEX gene expression tables for non-reproductive organs or in a collection *de novo* assembled transcriptomes from diverse healthy organs (see Methods). Reproductive organs, such as testis, were not included as this is an immunocompromised tissue that can also express antigens of interest for anticancer vaccination. Notably, we found that among tumor-specific transcripts lncRNAs and novel transcripts were more numerous than protein-coding genes (Figure 3a, Figure S8, Table S7 and S8). This was in sharp contrast with the observations for overall tumor expression, which was dominated by protein-coding transcripts (Figure 1f).

**Figure 3.**
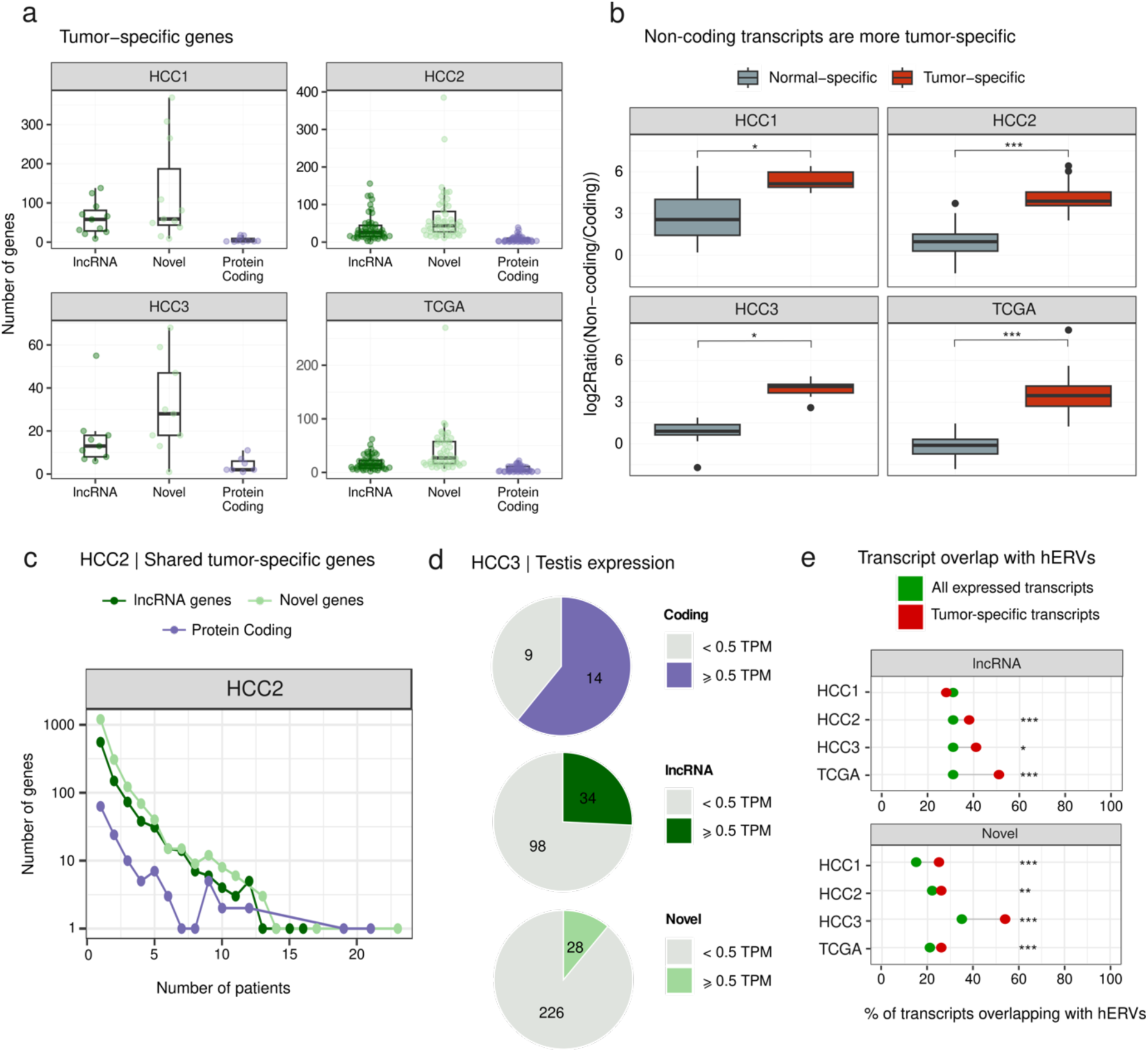
Most tumor-specific transcripts are non-coding. a. LncRNAs and novel transcripts tend to be more tumor-specific than coding genes. The number of different types of transcripts per patient and cohort is shown. **b. Tumor-specific versus normal-specific gene expression.** By paired Wilcoxon signed-rank test, we confirmed that the tumors are enriched in non-canonical genes with respect to coding ones. (HCC1 p-value = 4.883e-03; HCC2 p-value = 7.773e-10; HCC3 p-value = 7.813e-03; TCGA p-value = 2.469e-09) **c. Shared tumor-specific transcripts.** Despite the privacy of most tumor-specific transcripts, a subset are found in several patients. **d. Differences in the degree of expression in testis**. Proportion of tumor-specific transcripts that are also expressed in testis for different transcript types and datasets. **e. Proportion of lncRNA and novel transcripts overlapping with hERVs.** Differences between the fraction of lncRNAs overlapping hERVs in the complete transcriptome and in the tumor-specific transcriptome. Differences are significant in all cohorts except for lncRNA-HCC1 (pvalue < 0.05, Fisher test). Statistical significance is indicated as follows: ***p<0.001, * p<0.05.

One possibility was that the enrichment of lncRNAs and novel transcripts in the tumor-specific transcriptome could be attributed to the differences in expression levels or other factors unrelated to cancer. To assess this, we obtained the normal-specific transcriptome from each patient RNA-seq sample, by subtracting the transcripts that were also expressed in the tumor. We did not find a consistent enrichment of non-coding transcripts in this transcriptome (Figure S9). As a result, the ratio between tumor-specific and normal-specific transcripts was significantly higher in non-coding transcripts than in coding ones (Figure 3b). These results reinforced the notion that non-coding transcripts tend to be expressed in a tumor-specific manner much more frequently than coding ones.

Patient-shared tumor-specific antigens are of particular interest to develop therapies targeted at multiple patients. We inspected the number of patient tumor samples expressing the different types of tumor-specific transcripts. We found that the three types of transcripts – lncRNA, protein-coding and novel - could be shared by a significant fraction of the patients (Figure 3c, Figure S10, Table S9). No single tumor-specific transcript was shared across all patients; this was in line with previous reports for cancer/testis antigens, typically found in a percentage of the patient tumor samples ^36^.

Because many cancer antigens, such as the melanoma antigen genes (MAGE) are expressed in germinal cells, we wanted to investigate whether the same was true for tumor-specific non-coding transcripts. The analysis of the expression of tumor-specific protein-coding genes in testis confirmed that the majority of them showed significant expression in this organ (61-68% depending on the cohort). We also found that a subset fraction of the tumor-specific lncRNAs was expressed in testis (23-40%)(Figure 3d, Figure S11). In contrast, almost no expression in testis samples was detected for the novel transcripts.

LncRNAs are known to be enriched in remnants of transposable elements (TEs) when compared to protein-coding genes ^37^. We investigated if tumor-specific lncRNAs were different from the rest of lncRNAs regarding their TE composition, by inspecting their overlap with TE annotations derived from RepeatMasker^38^. No differences were found for most TE families except for human endogenous retroviruses (HERVs), that were significantly enriched among tumor-specific transcripts (Figure 3e, Table S10). HERVs have been reported to become activated during cancer ^39^. The demethylation of regions containing HERVs might play a role in the increased expression of HERV-containing lncRNAs.

### Tumor-specific antigens derived from ncORFs

It is currently unknown which is the relative contribution of ncORFs to the generation of tumor antigens when compared to other antigen sources. In order to estimate this, we first used NetMHCpan to predict all possible 9-mer peptides with strong binding affinity to MHC I (IC_50_ < 50 nM), for all tumor-specific ncORFs and coding sequences, as well as for peptides containing somatic mutations. The latter were obtained by performing variant calling directly from the transcriptomics sequencing data (HCC1, HCC2, HCC3) or from already available mutation data (TCGA). Since a peptide’s HLA binding affinity depends on the specific HLA allele, we inferred the HLA alleles of each patient using the RNA-Seq data and then performed predictions by HLA type. The proportion of strong binders (IC_50_ < 50 nM) among all possible peptides was similar for the different types of putative identified cancer-specific antigens (8.7%, 8.6%, 9.7% for lncRNA, novel and protein-coding, respectively) and slightly lower for mutated peptides (6.5%). Putative MHC I binders derived from mutations were essentially private (99.4%)(Figure 4a, Table S11 and Table S12). In contrast, an important fraction of the other tumor-specific antigens were shared across different patients. In the case of protein-coding genes (cancer/testis antigens) the fraction of predicted MHC I binders present in more than one patient was 42.6%. The equivalent figure for lncRNAs and novel transcripts was 28.4% and 16.2%, respectively.

**Figure 4.**
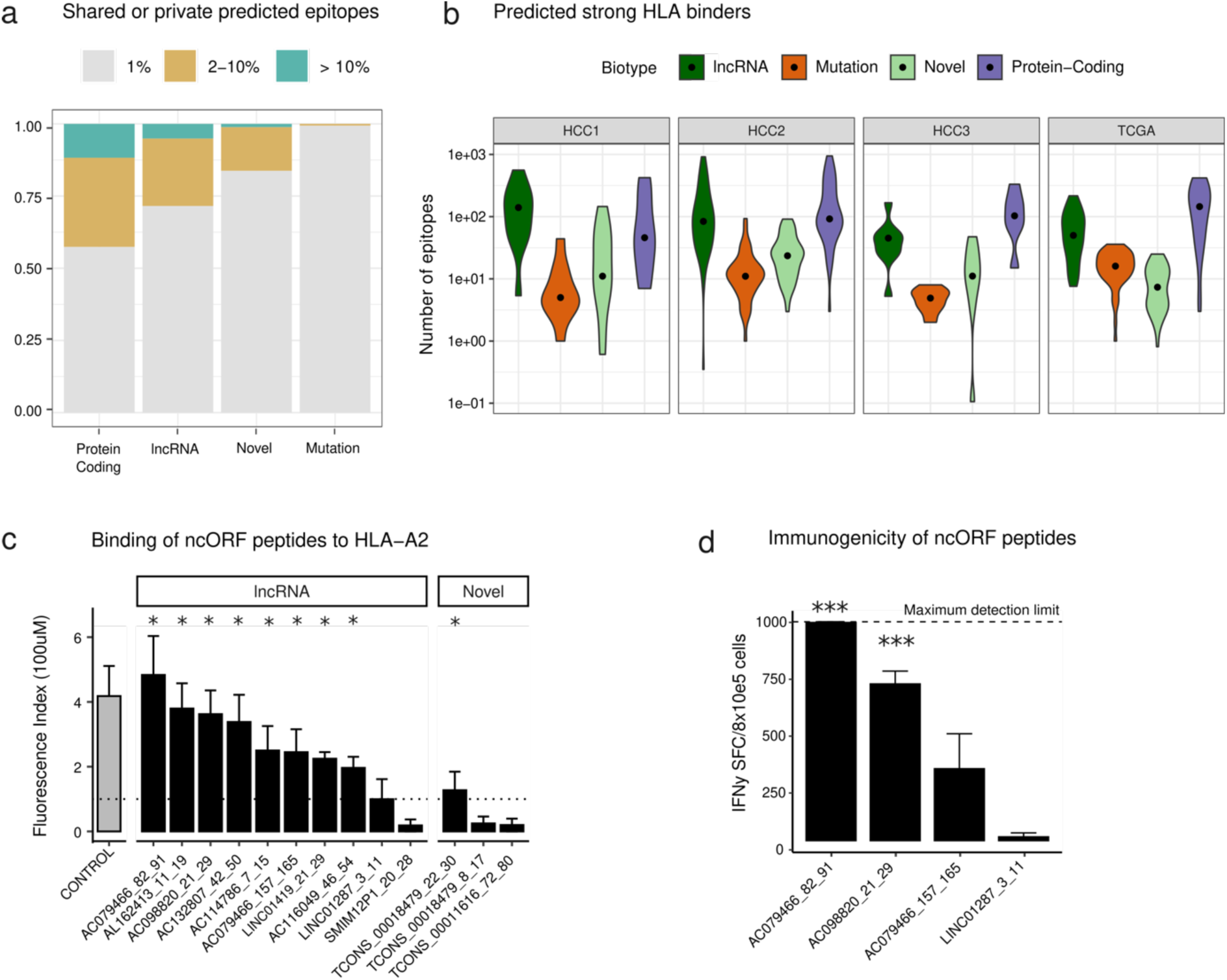
ncORFs make a substantial contribution to the HCC antigen landscape. a. Proportion of shared or private predicted antigens with respect to the total. Antigens derived from mutations are almost all patient-specific, whereas antigens derived from tumor-specific transcripts can be shared across patients. **b. Predicted number of antigens per patient and dataset**. Antigen load was predicted by selecting peptides with HLA-binding affinity IC_50_ < 50 nM as predicted by NetMHCpan, using patient-specific HLA allele information. For lncRNAs and novel transcripts it was then corrected by the translation index, which is the fraction of ncORF estimated to be translated by the analysis of Ribo-Seq data. **c. HLA-A*02.02 binding assays for ncORFs**. Binding affinity expressed as Fluorescence Index (FI) ± standard error of mean (SEM) for each peptide in an *in vitro* T2 cell binding assay. The FI value shown corresponds to the mean of two different assays (with two replicates each). A line at FI=1 indicates the expectation under no binding. We show the results of a Wilcoxon-Mann-Whitney test comparing the values obtained with the peptide and without the peptide (* p-value <0.05). Shown is the name of the transcript and the corresponding peptide start and end positions. According to this assay, 8 out of 10 ncORFs in lncRNAs and 1 out of 3 ncORFs in novel transcripts showed evidence of HLA binding. **d. IFN-γ ELISPOT assays.** The spleens of immunized mice were processed to measure the number of IFN-γ secreting cells (IFNγSFC). In the case of the first inspected peptide encoded by lncRNA AC07946, the measurement was > 1000 IFNγSFC/8e5 cells in all four replicates of the experiment. Differences with the background were computed and were significant in two cases (pvalue < 0.05, paired two-sample t-test). Of the four peptides tested, two yielded highly significant results (*** p-value <0.001, t-test), and the other two were not significant.

But, what do these observations imply at the level of individual patients? As we wanted to focus on non-self-peptides, we first identified and discarded any predicted strong binders matching other proteins/ORFs that were not tumor-specific. This eliminated 20-30% of the strong binders located in protein-coding genes but only 1-2% of those in lncRNAs and novel transcripts (Table S13). This means that, in general, peptides derived from ncORFs should not generate toxicities due to the cross-reaction with other proteins if used as vaccines. Next we multiplied the initial number of predicted strong binders in each of the patients by the translation index (0.116 for lncRNAs and 0.005 for novel transcripts). This index represents the overall fraction of the ncORF sequence space that is expected to be translated according to our previous analysis of Ribo-Seq data. The two most important contributors to the generation of antigens were tumor-specific lncRNAs and protein coding genes (median n=45-140 for lncRNAs and n=43-106 for protein-coding genes, depending on the dataset, Figure 4b and Table S14). In contrast, mutated peptides generated a relatively small number of predicted antigens (median n=5-16, depending on the dataset). In the case of novel transcripts, the low rate of translation meant that the estimates of the number of generated antigens were also low (median n=7-23, depending on the dataset). The average relative contribution of the different types to tumor-specific antigens, considering all 117 patients, was 49% protein-coding genes, 39% lncRNAs, 7% novel transcripts and 5% mutations.

### Experimental validation of predicted MHC I binders

In order to assess the reliability of the HLA binding predictions we selected 13 ncORF-derived peptides with high binding affinity for HLA-A*02.01 for *in vitro* testing using HLA-A*0.2.01+ T2 cells. Seven of them were shared by more than 10% of the patients and the rest by a smaller subset of patients (Table S15). The assays measured the peptide’s ability to bind to and stabilize MHC class I molecules, which are otherwise rapidly degraded in its normal peptide-unbound form.

We found that 8 out of 10 peptides derived from ncORFs encoded by lncRNAs, and 1 out of 3 peptides derived from ncORFs encoded by novel transcripts showed significant binding to the HLA molecule (measured as Fluorescence Index in Figure 4c)(Table S16). Overall, 9 out of 13 tested computational predictions were experimentally validated, which provides an estimate of around 70% of the predictions being actual HLA binders.

Four of the peptides with high predicted HLA-A*02.01 binding affinity were injected in HDD-DR1 mice, which contains genes encoding HLA-A*02.02 and HLA-DRB1*01, to test whether they could elicit an immune response (Figure 4d)(Table S17). Mice were immunized twice with the peptides and, fourteen days after the initial immunization, T cell response was measured through IFN-γ ELISPOT assay (Figure 4D). Peptide immunization resulted in strong IFN-γ signal in two cases, indicating T cell reactivity.

### Shared tumor-specific lncRNAs are expressed in different groups of patients

The analysis of the distribution of tumor-specific antigens across patient tumor samples is important to determine if they tend to cluster in the same group of patients or if they instead show a sparse distribution. If the second is true, multi-peptide vaccines could be effective in a larger number of patients. To examine this question, we selected transcripts shared by more than 10% of the patients that were expressed at relatively high levels (> 5 FPKM in at least one sample) and showed high tumor-specificity (expressed in less than 1% of the tumor adjacent samples considering all 117 patients)(Table S18). This resulted in 14 protein-coding genes and 33 lncRNAs (Figure 5a). Analysis of RNA-seq data from an unrelated cohort of 161 HCC tumor samples ^40^ showed that the majority of them (68%) were also expressed in more than 10% of the patients in the independent cohort (Figure S12). We also tested the expression of these transcripts in the thymus using previously published data ^16,41,42^. Expression in this organ induces central immune tolerance to self-peptides. No significant expression in thymic cells was observed for any of the transcripts, except for LINC02315 (Table S19).

**Figure 5.**
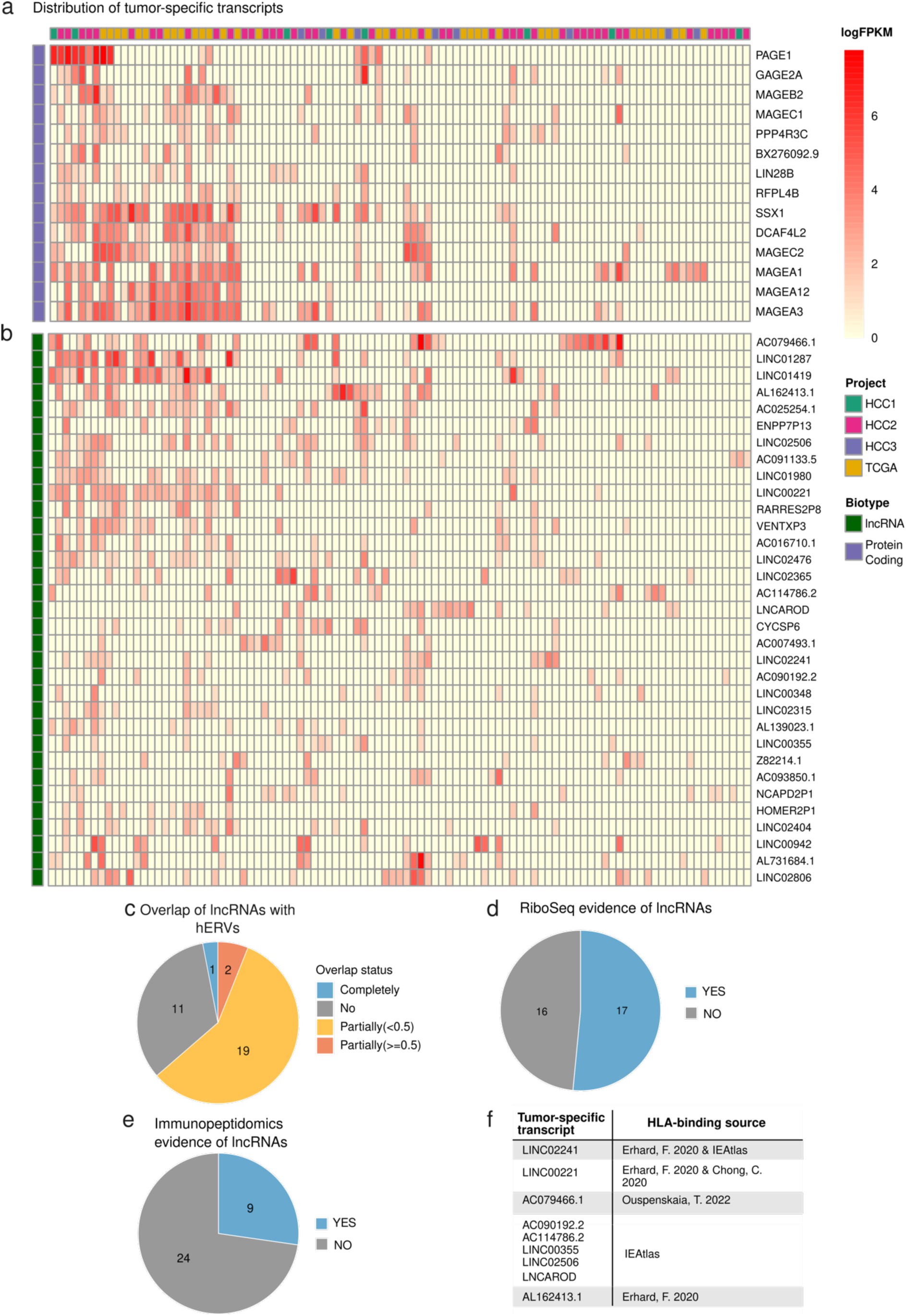
Distribution of shared tumor-specific transcripts in a large HCC cohort. Distribution of transcripts across different patients and datasets. Only genes which were tumor-specific in > 10% of the HCC tumor samples, expressed at more than 5 FPKM in at least one sample, and expressed in less than 1% of the normal liver samples (FPKM cutoff = 1), were considered. The intensity of the color in the cell reflects the level of expression (minimum of 1 FPKM). A. Distribution of protein-coding transcripts. Number of transcripts: 14. The transcripts tend to cluster in the left-most group of patients. B. Distribution of lncRNAs. Number of transcripts: 33. The transcripts are distributed in different groups of patients. C. Overlap with hERVs for this set of lncRNAs. Overlap is based on genomic coordinates. D. Proportion of lncRNAs with detected translated ncORFs. Prediction of ncORF translation was performed using Ribo-Seq data from a different HCC cohort of 10 patients (HCC4). E. Proportion of lncRNAs containing ncORFs with immunopeptidomics evidence in other cancer datasets. Details of the lncRNAs and the works that reported HLA binders are shown.

The majority of protein-coding genes in this set tended to cluster in the same group of patients, suggesting similar gene expression activation mechanisms (Figure 5a). Instead, lncRNAs tended to be more scattered across patients (Figure 5b). Accordingly, gene expression correlation values between pairs of protein-coding genes tended to be higher than between pairs of lncRNAs (Figure S13). In particular, the average correlation between any two protein-coding genes was 0.44, whereas for lncRNAs it was 0.19. Furthermore, most protein-coding genes (11 out of the 14) were located in the X chromosome, whereas lncRNAs were dispersed across different chromosomes (Table S20). In line with the previous observation that tumor-specific lncRNAs were enriched in HERV-derived sequences, we found that 21 out of the 33 lncRNAs showed some degree of overlap with HERVs (Figure 5c)(Table S21). This points to a possible link between suppression of the silencing of HERV-containing genomic regions in tumors and the activation of new transcripts.

### LncRNAs are frequently translated and produced HLA-bound peptides

We detected signatures of translation in more than half the lncRNAs in this subset (17 out of 33), using Ribo-Seq data from the HCC4 cohort and the same procedure as previously described (Figure 5d)(Table S21). In addition, 9 lncRNAs encoded HLA-bound peptides according to the results of different immunopeptidomics studies^17,18,20,43^(Figure 5e and 5f)(Table S21). Because these studies did not include HCC tumors, this indicates that the lncRNAs are expressed and translated in other cancer types as well. These results support that lncRNAs are a frequent source of shared tumor-specific antigens.

There were 10 lncRNAs that were expressed in more than 20% of the tumor samples (LINC01980, AC025254.1, LINC02806, LINC02476, LINC02506, AL162413.1, LINC00221, LINC01287, AC079466.1 and LINC01419), compared to 5 protein-coding genes (MAGEA1, MAGEA3, SSX1, DCAF4L2, MAGEC2). We found evidence of translation for 7 out of the 10 lncRNAs (Table S21). In addition, three of these lncRNAs had immunopeptidomics evidence (AL162413.1, LINC00221, AC079466.1). The lncRNA LINC00221 has previously been shown to have an anti-tumor effect by inhibiting HCC cell growth, migration and invasion^44^. In contrast, AC079466.1, has been associated with poor prognosis in HCC ^45^. One of the peptides encoded by this protein showed high HLA binding affinity in the experiments with T2 cells as well as the ability to induce high interferon production in mice (Figure 4d).

## DISCUSSION

In this work we have shown that peptide presentation from ncORFs significantly expands the number of tumor-specific antigens over traditional sources, providing additional targets for immunotherapy. We have designed a pipeline that integrates different sources of data from tumors and matched healthy tissue to characterize the tumor-antigen landscape, which we have then applied to a meta-cohort of 117 HCC patients. This pipeline successfully identifies known cancer/testis antigens as relevant in hepatocellular carcinoma, including the melanoma-associated antigen (MAGE) family, cancer-testis antigen G antigens (*GAGE*) or the synovial sarcoma X breakpoint 1 antigen (SSX1)^5,36^. But it also identifies ncORFs in lncRNAs as important players in defining tumor antigen landscape. In contrast to canonical cancer antigens, ncORFs do not show a strong clustering in the same subset of patients. This means that they could potentially be used to target a larger number of patients.

We studied both lncRNAs and non-annotated, novel, transcripts in the tumor as a possible source of tumor antigens. Using ribosome profiling data from HCC patients, we found that translated ncORFs concentrated in lncRNAs, whereas almost no translation signatures were detected in novel transcripts. Ribosome profiling has been instrumental to uncover the translation of many small ORFs that had not been initially annotated as coding ^15,31,46^. These studies have also revealed that ncORFs in lncRNAs and mRNA untranslated regions (UTRs) can initiate translation from codons other than ATG ^34^. Here we leverage HCC Ribo-Seq data to infer the rate of translation of the total ncORFome in HC tumors, as well as to detect translated ncORFs in a subset of patient-shared tumor-specific lncRNAs. Our results are in line with the notion that many lncRNAs contain translated ncORFs. The median length of the translated ORFs was 39 amino acids, similar to that described in a large set of Ribo-seq ORFs recently catalogued in GENCODE (average 44 amino acids)^15^. We also gathered evidence that ncORFs can generate in HLA-bound peptides by querying existing tumor immunopeptidomics data. Taken together the data showed that peptides derived ncORFs can be highly tumor-specific, shared across patients and presented to HLA receptors.

Previous studies on cancer lncRNAs have mainly focused on lncRNA tumor overexpression ^47,48^. It has been shown that some of the over-expressed lncRNAs have oncogenic activities, such as promoting cellular proliferation and interfering with epithelial-to-mesenchymal transition regulators ^48^. The activity can be mediated by the RNA molecule itself or by encoded microproteins. One example of the latter class is LINC00998. This lncRNA codes for the SMIM30 micropeptide, which activates the MAPK pathway ^49^. In our study we instead focused on lncRNAs that are expressed in a tumor-specific manner, as they are likely to be more relevant for cancer immunotherapy.

Although the functions of the majority of the identified tumor-specific lncRNAs remain uncharacterized, several of them have previously been suggested to have roles in cancer. One example is LINC02241, which we found expressed in 13.6% of the HCC samples, and which has been associated with poor prognosis in colorectal cancer ^50^. Another example was AL162413.1 (22.2% of HCC samples), described as a potential biomarker of oral squamous cell carcinoma ^51^. Lastly, LNCAROD, found in 14.5% of HCC samples, has been associated with attenuation of cell proliferation, whereas the opposite effect is observed when the gene is overexpressed ^52^. These three lncRNAs, together with another six ones, had been reported to encode HLA-bound peptides in other cancer cell types. This highlights their pervasiveness in tumors of different origin and their potential utility for pan-cancer therapies.

Epigenetic dysregulation of the cancer genome leads to the expression of genomic regions that are generally silent ^53^. For example, it was shown that the promoter of the polycistronic transcript meloe, which encodes the MELOE-1 and MELOE-2 cancer-specific peptides, is hypomethylated in melanoma^54^. HERV sequences cover about 8% of the genome and can become reactivated in cancer ^55^. We found an enrichment of HERV sequences among tumor-specific transcripts, suggesting that they might facilitate the expression of cancer-specific lncRNAs. Interestingly, HERV signatures have been shown to predict immunotherapy response in clear cell renal cell carcinoma ^56^, suggesting a link between transcriptional activation of HERV-containing regions and tumor antigen production. Changes in transcription factor activity or even the formation of novel transcription factors, could also have an effect. One known case is the formation of a chimeric transcription factor in Ewing sarcoma, which leads to the expression of a new set of spliced and polyadenylated transcripts, some of which can translate new peptides ^57^.

For tumor antigens to be of therapeutic usefulness, they must be capable of triggering an immune response against the tumor. MAGEA1 was the first of a series of cancer/testis antigens which have been shown to be recognized by cytolytic T lymphocytes ^58^. Several of these antigens have since then been used to develop vaccines, with some having shown clinical benefits ^7^. However, the clinical trials have also revealed that caution needs to be taken to avoid toxicities, for example induced by the cross-reactivity between several MAGE-A family members, not all of which are expressed in a tumor-specific manner ^59^. These problems are not expected for ncORFs, as lncRNAs do not cluster into families ^30^. Consistently, we found nearly no matches between predicted strong binders encoded by lncRNAs and non-tumor-specific proteins, whereas the same was not true for cancer/testis antigens. We observed that, similar to canonical cancer antigens, lncRNAs showed expression in some, but not all, patients. The difference was that they were more diverse in their distribution patterns, which means that they could potentially be used to target groups of patients that would not be targetable with classical cancer/testis antigens.

We tested four ncORF-derived peptides in mice transgenic for HLA-A*02.01, and found that two of them could generate a significant immune response. The immunogenicity of this class of peptides in humans remains largely to be investigated. One previously described case is MELOE-1, involved in T cell transfer efficiency ^8^. Other examples include peptides arising from the tumor-specific expression of intronic regions or alternative frames ^60,61^. More recently, noncanonical splicing junctions between exons and TEs have also been reported to be a source of immunogenic antigens in cancer by stimulating human T cell populations ^62^.

In the study we combined data from different sources, which poses some limitations. For example, we used Ribo-Seq data from a different HCC cohort than the RNA-Seq data, and this decreased our ability to detect translation in transcripts that were highly patient-specific. We also used immunopeptidomics data from cancer types other than HCC to further support HLA binding of ncORF-derived peptides. In future studies, using mass spectrometry data from the same tumors/matched tissue should provide more accurate estimations of the number of HLA-bound peptides that are tumor-specific. At the same time, it would allow testing if ncORFs, which encode proteins that are potentially more unstable, result in a larger number of HLA-bound peptides than other types of proteins ^21^. Finally, testing immunogenicity in T cells from patients expressing the ncORF candidates would be a necessary step to guide any future vaccine development.

In summary, our study has investigated the prevalence of ncORFs in a composite large cohort of tumor/matched HCC samples, revealing that ncORF-derived peptides can be highly tumor-specific, patient-shared and presented by HLA receptors. The analysis has identified several promising candidates that might be involved in tumorigenesis and/or be capable of activating T-cell responses after vaccination or suppression of checkpoint inhibitors. This study encourages research of ncORFs in other cancer-types and open new avenues for treatment.

## METHODS

### Pre-processing of raw sequencing data

We obtained RNA-Seq data containing HCC tumor/normal paired data from GEO entries GSE101432 ^24^, GSE77314 ^63^, GSE193567 ^25^, GSE112705 ^26^, as well as from TCGA (Cancer Genome Atlas Research Network 2017). In the case of GSE112705 we also downloaded and analyzed Ribo-Seq data. These datasets were named HCC1, HCC2, HCC3, HCC4 and TCGA, respectively. The sequencing reads were for total RNA except for TCGA, which was polyA+ RNA. Files of raw reads were downloaded from the Gene Expression Omnibus (GEO) database and the SRA Toolkit (v 2.9.2)(https://trace.ncbi.nlm.nih.gov/Traces/sra/sra.cgi?view=software) was used to convert SRA to FASTQ format. For TCGA data, files with mapped reads (bam files) were downloaded from the GDC Data Portal ^64^ and reverted to FASTQ format using SamToFastq from Picard Toolkit (v 2.25.1) (http://broadinstitute.github.io/picard/). RNAseq reads were quality assessed using both FastQC (v0.11.5) and FastQScreen (v0.14.0) software ^65^. All selected samples passed the quality control. The Cutadapt (v.2.1) program ^66^ was used to trim 3’ adapters from the raw reads with -O 5 -q 30 -m 26 parameters. Sequencing reads were aligned to the human reference genome GRCh38/p13 using two-pass alignment with STAR (v2.7.1)^67^ to improve the quantification of novel splice junctions. Only uniquely mapped reads were considered.

### Novel transcript assembly

We assembled the transcriptome of each tumor/normal sample separately with StringTie (v2.0)^68^ in a conservative mode, using GENCODE annotation version 38 as the reference annotation file. In order to recover novel, non-annotated transcripts, we extracted the set of non-annotated transcripts and selected those that did not overlap with any transcript in the human reference annotation using BEDTools (v2.2.1)^69^. The assembly of novel genes worked well in all cohorts except for one patient sample from HCC1, which appeared to have an abnormally high number of novel transcripts. This outlier was removed from further analysis.

In datasets HCC1 and HCC3, with strand-specific RNA-Seq data, novel transcripts had a known orientation (mapped to the plus or minus genomic strand) and a similar number of exons distribution than annotated lncRNAs. In contrast, in datasets HCC2 and TCGA, which were not strand-specific, the vast majority of the novel transcripts corresponded to single exon genes, and their orientation was not known. In this case we imputed the orientation from miTranscriptome transcripts with overlapping genomic coordinates. If no matches in miTranscriptome could be found, the transcripts were discarded. Neither novel transcripts shorter than 300 pb nor those longer than the longest annotated tumor lncRNA (*KCNQ1OT1, 91666 nucleotides*) were considered for further analysis.

### Transcript expression quantification

For each patient we built a complete transcriptome by merging the annotated genes with the novel transcripts obtained by *de novo* transcript assembly and not matching any annotated genes. In the case of annotated genes we considered coding and long non-coding RNAs (lncRNAs), and kept the longest transcript per gen. LncRNAs included the class “processed pseudogenes”. To quantify gene expression we used featureCounts ^70^, from the Subread package (v2.0.3), in stranded mode whenever possible. Next, we converted the counts (uniquely mapped reads) to fragments per kilobase per million mapped reads (FPKM). Transcripts with expression values lower than 1 FPKM were not considered for downstream analysis. In the case of HCC1 dataset this cut-off was increased to 2 FPKM because lncRNAs/novel transcripts tended to have higher expression values in general. Using the 2 FPKM cut-off ensured that a similar proportion of lncRNAs/novel transcripts were recovered in this dataset when compared to the other ones.

### Merging novel transcripts from different patients

Gffcompare software ^71^ was used to merge novel transcripts from different samples on the basis of overlapping genomic coordinates and obtained a non-redundant set of representative transcripts. This step was necessary because the same transcript might be reconstructed in slightly different ways in different samples, resulting in different but overlapping genomic coordinates. Each representative transcript had a unique identifier, which we used to track the transcript across patients. We run BLASTN (v 2.11)^72^ with default parameters to detect possible homology between the representative transcripts and ribosomal RNA genes. Those that had significant sequence homology with rRNA genes were discarded (E-value < 10^-3^).

### Prediction of translated open reading frames

We defined ncORFs as starting with an ATG, ACG, CTG, GTG or TTG and ending with a stop codon. The ncORFs had a minimum length of 30 nucleotides. When two ncORFs overlapped in the same frame we selected the longest one. For protein-coding genes only the annotated coding sequence was considered. Translation was predicted using RibORF (v. 1.0)^13^, which generates a score by combining the three nucleotide periodicity and homogeneity of the signal along the ORF. In order to increase sensitivity we first merged the reads from 10 ribosome profiling tumor samples (HCC4). We obtained a total of 99 Million mapped Ribo-Seq reads. The reads had an average periodicity (proportion of reads in the correct frame) over 0.5 (e.g. 0.58 and 0.51 for 28 bp and 29 bp sequences, respectively). A minimum of 5 footprints and a RibORF score of at least 0.5 were the criteria used to classify any ncORF as translated. From the total set of putatively translated ncORFs, we eliminated those overlapping in the same frame, keeping only the longest one. We ended up with a non-redundant list of translated ncORFs per dataset.

Many of the lncRNAs and novel transcripts detected in the 117 HCC tumor samples are patient-specific or restricted to a few patients. Because the Ribo-Seq data is for a different cohort of 10 HCC samples, we could expect low sensitivity. In order to obtain reliable estimates of the level of translation of lncRNAs and novel transcripts we focused on those expressed in at least 90% of the patients in the cohort of interest as well as in the cohort with Ribo-Seq data. In the latter cohort expression was determined using the available RNA-Seq data and a cut-off of 1 FPKM. We also used the Ribo-Seq data to investigate the translation of a subset of tumor-specific transcripts expressed in more than 10% of the individuals of the 117 HCC meta-cohort.

### Tumor-specific gene subset

An expression cutoff value of > 1 and < 0.1 FPKM in tumor and adjacent normal samples, respectively, was established to select transcripts expressed only in the tumor sample in each of the patients. In the case of the HCC1 dataset the cut-off was >2 and < 0.2 FPKM, respectively, due to overall higher expression levels of non-coding transcripts (Figure S1). We also collected expression data from the Genotype-Tissue Expression (GTEx) project ^73^, which includes RNA-Seq experiments from a wide spectrum of body tissues. The data was used to discard genes with a median expression higher than 0.5 TPM (transcripts per Million) in any non-reproductive tissue. Expression in testis and/or ovary was not considered because germinal cells do not express MHC receptors and thus no antigens can be detected by the immune system. In the case of novel transcripts, which were not represented in GTEX, we used *de novo* transcript reconstructions using publicly available RNA-Seq data from a range of human tissues (brain, cerebellum, heart, kidney, liver, testis)^74^(Table S22). Novel transcripts with a median expression higher than 0.5 TPM in this set of healthy tissues (with the exception of testis) were removed as well.

### Analysis of non-coding transcripts overlapping TE annotations

We examined the genomic overlap between lncRNA and novel transcripts on one hand, and TE sequences on the other. We sourced TE annotations from the UCSC Genome Browser’s RepeatMasker track for the GRCh38/hg38 genome assembly (http://genome.ucsc.edu/)^75^. Then, we removed low complexity regions, simple repeats, satellites, rRNA, scRNA, snRNA, srpRNA and tRNA to only keep TE instances. Moreover, we only kept TEs with known strand information (i.e. features with a strand different than “+” or “-” were discarded). In order to identify which tumor-specific lncRNA / novel transcripts were overlapped by TEs, we used the “findOverlaps” function from the “GenomicRanges” package ^76^, requiring a minimum overlap of 1 base pair on the same strand. Next, we computed the fraction of lncRNA/novel transcript length occupied by TEs. To do so, we accounted for the possibility of two or more TEs overlapping between them while contained in the lncRNA / novel transcript, avoiding counting this TE overlapping length twice. We then compared the frequencies of different classes of TE elements (LINE, SINE, Retrotransposons, hERV, RNA) between tumor-expressed and tumor-specific non-coding transcripts. Only hERVs showed a significant enrichment in the tumor-specific subset of transcripts that was consistent across HCC cohorts.

### Identification of potential MHC I bound peptides

We used optitype from the nf-core/hlatyping pipeline (https://nf-co.re/hlatyping) and arcasHLA ^77^ to determine patients’ 4-digit HLA I. Next, netMHCpan 4.1 ^78^ was used to predict the potential immunogenicity of the previously identified sequences with coding potential. We derived all possible 9-mer peptides from canonical protein sequences and non-canonical ORFs and selected those with predicted IC_50_ < 50 nM (concentration that inhibits 50% binding of the fluorescein-labeled reference peptide) as strong MHC I binders.

### Variant Calling

For HCC1, HCC2 and HCC3 datasets, we used GATK4 best practices ^79,80^ pipeline described for variant calling in somatic RNA-seq data with tumor and matched normal samples. Mutect2 ^81^ was used to detect SNVs that were later filtered according to standard quality metrics. For TCGA, we used previously define somatic mutations (Cancer Genome Atlas Research Network 2017). Only mutations from genes expressed in the patient sample were considered. In both conditions, we required a minimum of total depth of coverage > 10 and a minimum of three reads supporting the alternative variant to obtain high-confidence SNVs. The functional annotation of the identified somatic mutations was done with Ensembl Variant Effect Predictor tool (v.98)^82^ and only those whose consequence is missense (change of amino acid) were maintained. We identified potential neoantigens arising from the mutations in the same manner as for tumor-specific translated products, using a sliding window around the mutated aminoacid.

### HLA binding studies

Potential epitopes binding to HLA-A*02.01 were predicted *in silico* with NetMHCpan 4.1 ^78^ from 9- mers derived from ncORFs located in tumor-specific non-coding transcripts. The selected 9mers had a range of predicted HLA-A*02.01 affinities between 8.04 and 68.04 nM, and were found in 13 different lncRNAs and 3 novel transcripts. We prioritized lncRNAs found in a wide range of patients (9- 31 patients). For HLA-A*02.01 binding assays, peptides were synthesized with a purity ≥ 80% at GeneCust. HLA-A*02.01+ T2 cells were used to determine peptide binding to HLA-A*02.01 molecules.

Cells (2.5 × 10^5^/well) were cultured in 96-well microplates with 100 μM of the corresponding peptide and incubated overnight at 37 °C. Samples were then incubated with Beriglobin (800 μg/mL), stained with FITC-labeled anti-HLA-A*02 (*Genetex*) (2 mg/mL, 15 min at room temperature) and mean fluorescence intensity (MFI) was determined by flow cytometry. Influenza M protein was used as a positive control. Peptide binding was expressed as Fluorescence index (FI) using the following formula: (MFI with peptide – MFI without peptide)/ MFI without peptide. We performed two independent experiments for each peptide, each time taking two measurements.

### Immunization experiments

Eight-week-old female HHD-DR1 mice, transgenic for human HLA-A*02.01 molecules ^83^ were used. After study approval by the ethics review committee (ref #036-21), mice were bred and housed under pathogen-free conditions in the animal facility of the *Center for Applied Medical Research* (CIMA). Mice (n = 4/group) were immunized with peptides (100 nmoles/peptide), poly(I:C) (50 μg/mouse), and αCD40 (50 μg/mouse). The peptides and adjuvants were administered simultaneously via subcutaneous (s.c.) injection of 100 μL of the mixture resuspended in PBS. Mice received a boost on day 7 and were sacrificed on day 14.

### IFN-γ ELISPOT

The spleens of immunized mice were processed to measure the number of IFN-γ secreting cells. Splenocytes (8×10^5^ cells/well) were stimulated for 24 h with peptides (10 μg/ml). The number of IFN- γ secreting cells was quantified by ImmunoSpot automated counter (*Cellular Technology Limited*) using the Spot 3 CTL CellCounting software.

### Identification of frequently-occurring tumor-specific transcripts

We selected transcripts that were tumor-specific in at least 10% of the patients (12 or more out of 117), with an expression value higher than 5 FPKM in at least one patient, and with a very high overall tumor-specificity. The latter was defined as expression in less than 1% of the 117 normal adjacent normal samples using an expression cut-off of 1 FPKM for HCC2, HCC3 and TCGA or 2 FPKM for HCC1. We identified 14 protein-coding genes and 33 lncRNAs that met these requirements. To validate these findings, we downloaded the normalized expression data from an external dataset with 161 tumor samples from HCC patients from the International Cancer Genome Consortium (ICGC) data portal (http://dcc.icgc.org/; release 28). For the highly frequent tumor-specific genes, we measured the proportion of tumor samples that were expressing each gene. We used R (version 4.1.2) to measure the correlation between the percentage of patients expressing each candidate gene per study. We also analyzed the expression of this subset of tumor-specific transcripts in the thymus using 30 publicly available thymic epithelial cell (TEC) samples from two GEO entries: GSE127825 ^16,42^ and GSE201719 ^41^(Table S23). TEC samples were aligned to the genome and quantified using the abovementioned pipeline. Transcripts with a median expression higher than 0.5 FPKM were considered to be expressed in thymus.

### Statistical Analysis

Statistical analyses were performed using R (version 4.1.2). Comparisons between two distributions were performed using the paired Wilcoxon signed-rank test (Figure 1b, Figure 3b, Figure S8) or the Kolmogorov Smirnov test (Figure 2e). The difference between two proportions was assessed using Fisher exact tests (Figure 3e). After analyzing Gaussian distribution of data with the Shapiro test, Wilcoxon signed-rank test was performed to assess the differences between the mean fluorescence intensity (MFI) of the peptides with respect to the MFI of the peptide control (Figure 4c). To test the immunogenicity of the ncORF peptides compared with their background signal, we performed paired two-samples t-tests.

## SUPPLEMENTARY MATERIAL

Supplementary figures are in a supplementary file. Supplementary tables are in Figshare repository: https://doi.org/10.6084/m9.figshare.24448723.

## ACKNOWLEDGEMENTS

We gratefully thank José Carlos Montañés and Chris Papadopoulous for the useful insights into Ribo-Seq data analysis, as well as our research group for their continuous support throughout the research. The work was supported by the following grants and agencies: 1. Research projects PID2019- 105595GB, PID2021-122726NB-I00 and PID2021-128791OB-I00 funded by MCIN/AEI/10.13039/501100011033 and by “ERDF: A way of making Europe”, by the “European Union”; 2. PI22/00171, funded by Instituto de Salud Carlos III (ISCIII) and co-funded by the European Union; 3. 0011-1383-2019-000006 and 0011-1411-2021-000070, funded by Gobierno de Navarra; 4. RTI2018-101759-B-I00 and the Instituto de Salud Carlos III, which finances Centro de Investigación Biomédica en Red de Enfermedades Hepáticas y Digestivas (CIBEREhd) and Red Española de Terapias Avanzadas TERAV ISCIII (RICORS: (RD21/0017), financed by the EU (NextGenerationEU. Plan de Recuperación Transformación y Resiliencia). 5. AECC IDEAS20169FORT by Scientific Foundation of the Spanish Association Against Cancer; 6. 2021SGR00042 by Generalitat de Catalunya; 7. Ayudas Fundación BBVA a Proyectos de Investigación Científica en Biomedicina 2021; 8. European Reseach Council Advanced Grant NovoGenePop 101052538. Views and opinions expressed are however those of the authors only and do not necessarily reflect those of the European Union or the European Research Council. Neither the European Union nor the granting authority can be held responsible for them.

## AUTHOR CONTRIBUTIONS

M.E.C. compiled the samples and performed most computational analyses. P.T., M.T., P. F. and P.S. performed immunization experiments in mice. M.R. and P.S. performed in vitro HLA-peptide binding experiments. J.R-O. analyzed ribosome profiling data. B.C-S. and R.C. analyzed TEs overlapping tumor-expressed lncRNAs. M.E.C, J.R-O., P.S. P.F., J.P-B., M.M.A. contributed to the conceptualization of the study and design of experiments. P.F., J.P-B. and M.M.A supervised the study. M.E.C. and M.M.A wrote the manuscript with input from all authors.

