## Supplementary Figures S1-S13 for "Non-canonical ORFs are an important source of tumor-specific antigens in a liver cancer meta-cohort"

**Figure S1. Establishing a gene expression threshold for different HCC tumor RNA-Seq datasets**. Some of the lncRNAs/novel transcripts showed very low levels of expression, making them of little relevance with regards to antigen HLA presentation. For this reason, we established a minimum expression threshold, which we applied to all transcript types. In three of the four datasets (HCC2, HCC3 and TCGA) we used a cut-off of 1 FPKM (solid line), which approximately separates the distribution of protein-coding and non-coding genes. In dataset HCC1 we used a cut-off of 2 FPKM (dashed line) using the same criteria.

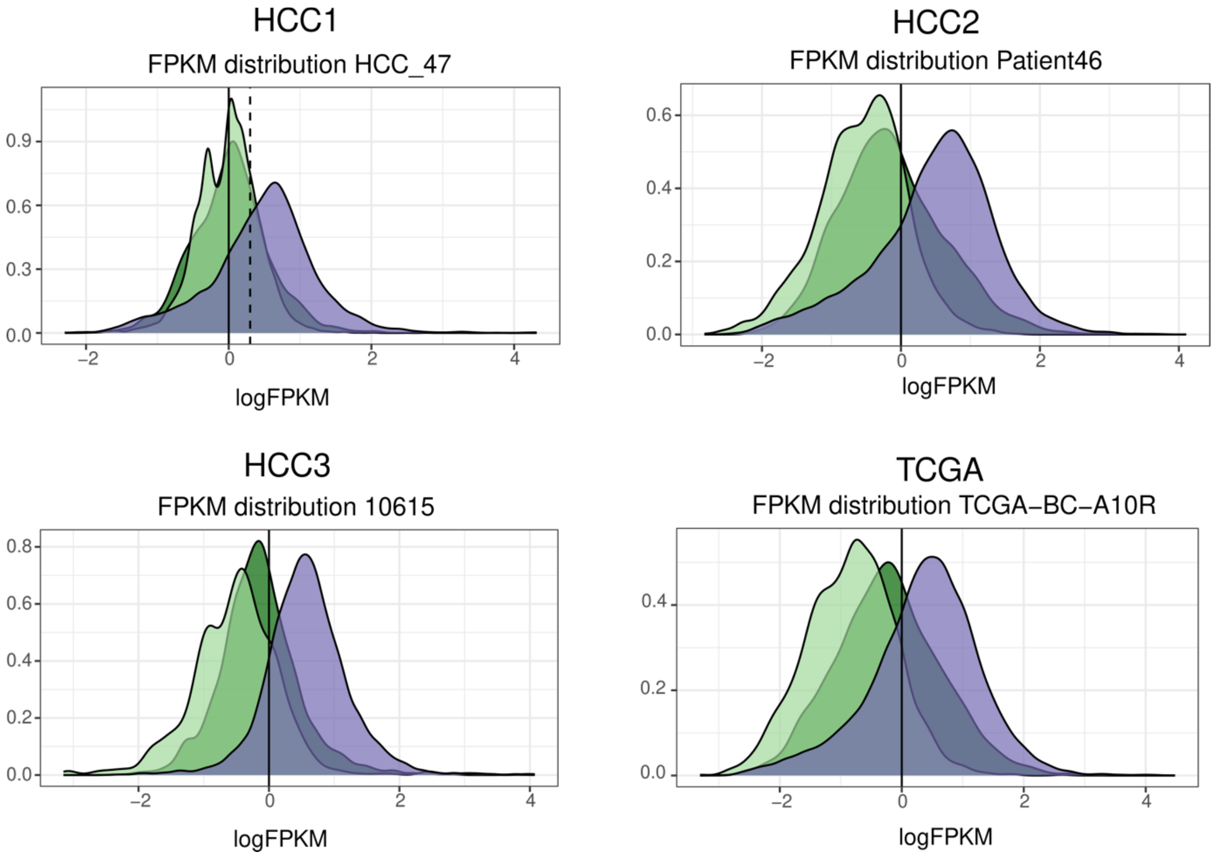

**Figure S2. Exon counts per dataset.** The average number of exons in the transcripts from the different tumor RNA-Seq datasets was calculated. As expected, lncRNAs and novel transcripts tend to have a lower number of introns when compared to coding genes. In unstranded RNA-Seq data the recovery of multiexonic novel transcripts is decreased with respect to stranded datasets. The equivalent Figure for dataset HCC3 is shown in the main manuscript file.

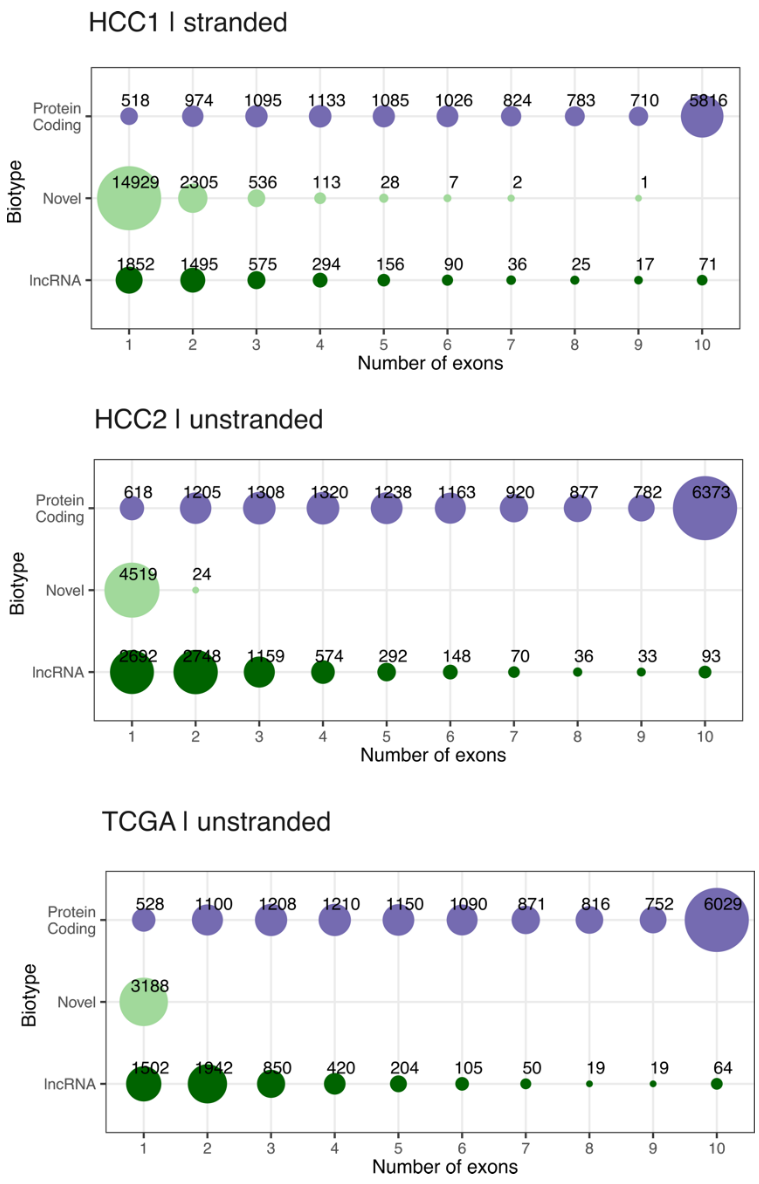

**Figure S3. Identification of novel transcripts in miTranscriptome.** Matches of novel transcripts assembled in this study and transcripts already defined in miTranscriptome were based on genomic coordinates overlap. The percentage of matched transcripts ranges from 57% to 99% depending on the dataset. Strand-specific datasets (HCC1 and HCC3) showed a smaller percentage of matches than non-stranded ones (HCC2 and TCGA).

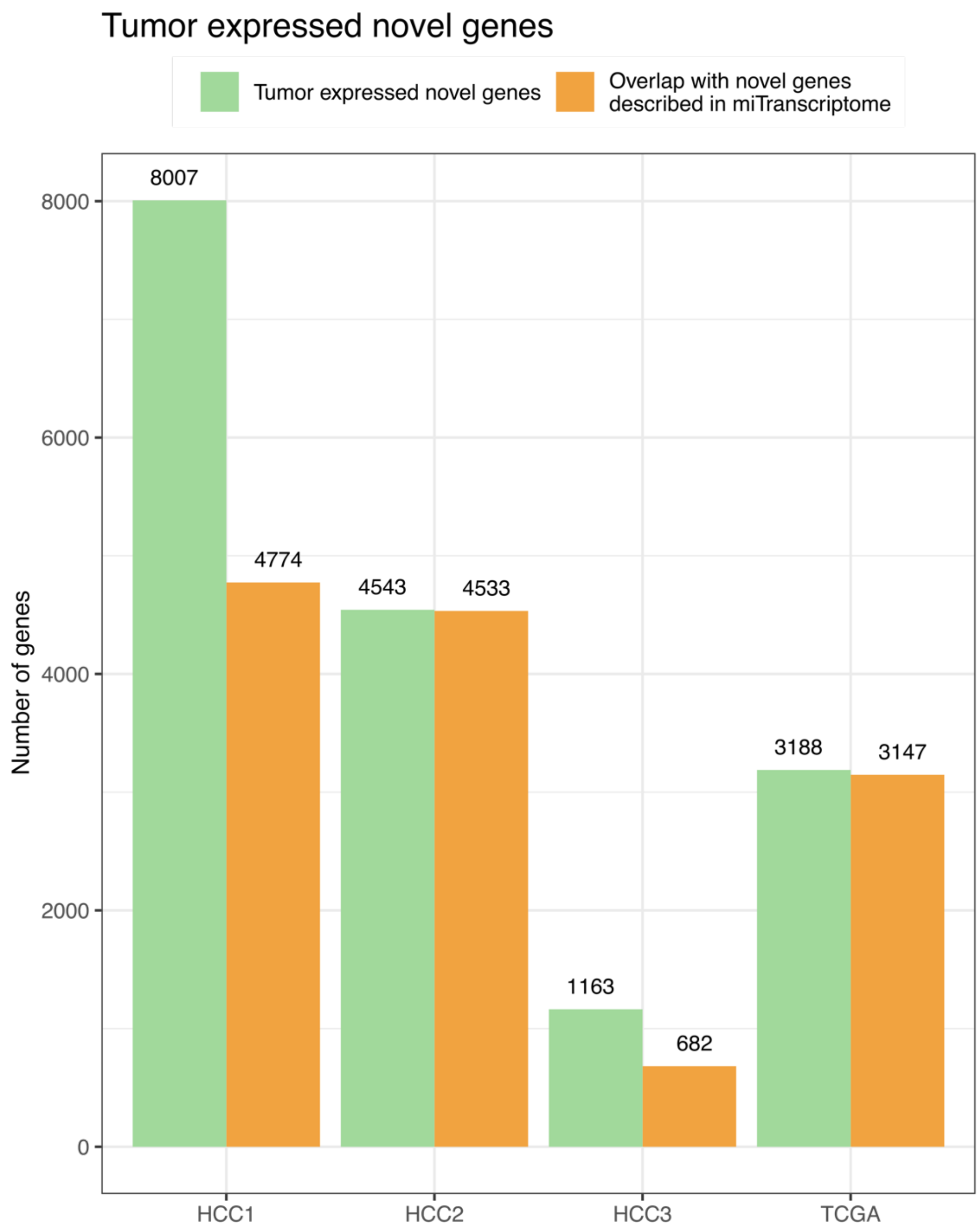

**Figure S4. The HCC transcriptome per patient sample and dataset.** Number of genes expressed (no expression cut-off). The longest transcript per gene is considered. Novel genes were reconstructed from the RNA-seq data.

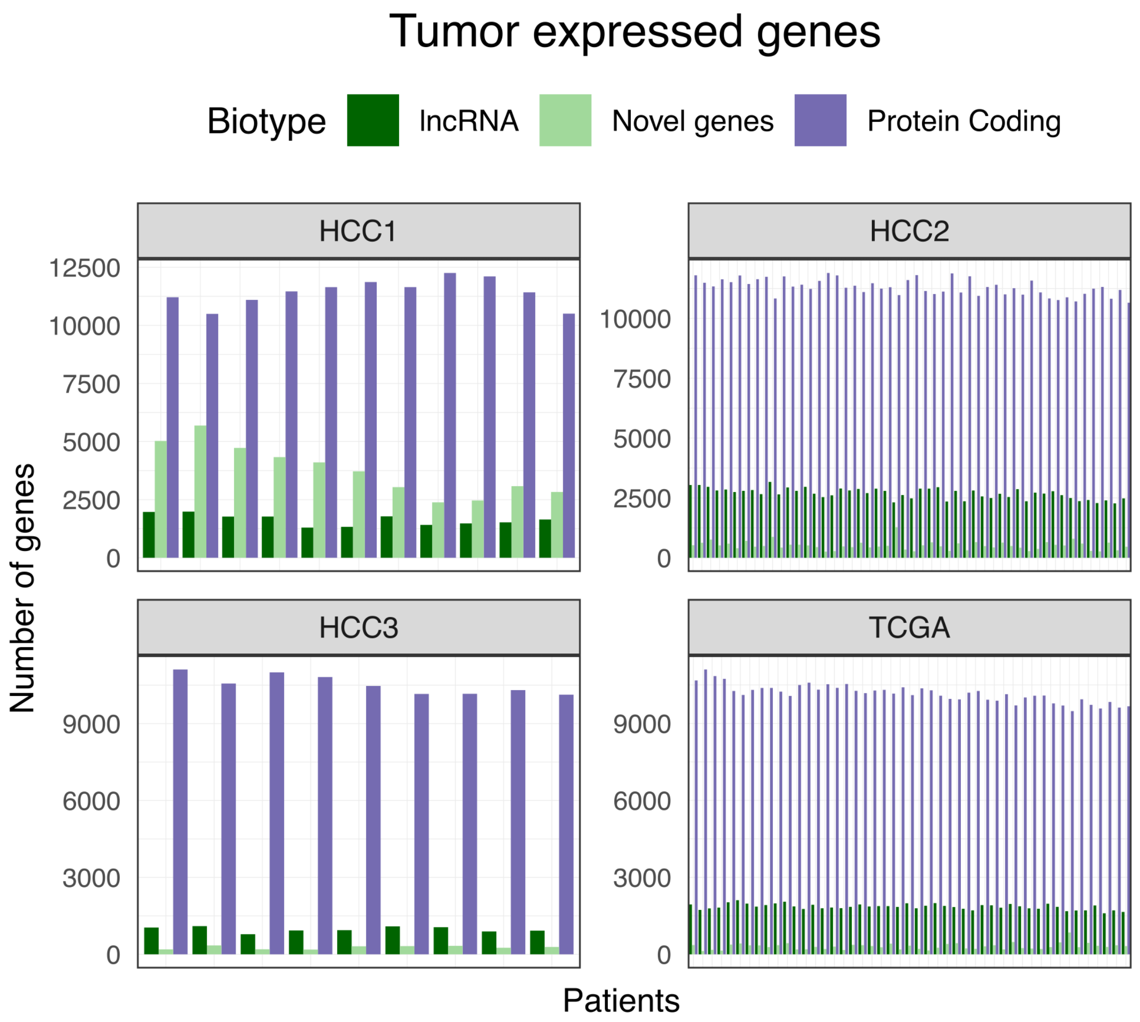

**Figure S5. Number of patients that share a given transcript.** When comparing the genes expressed in the tumors across patients, lncRNA are less shared than coding genes and novel transcripts were mostly private.

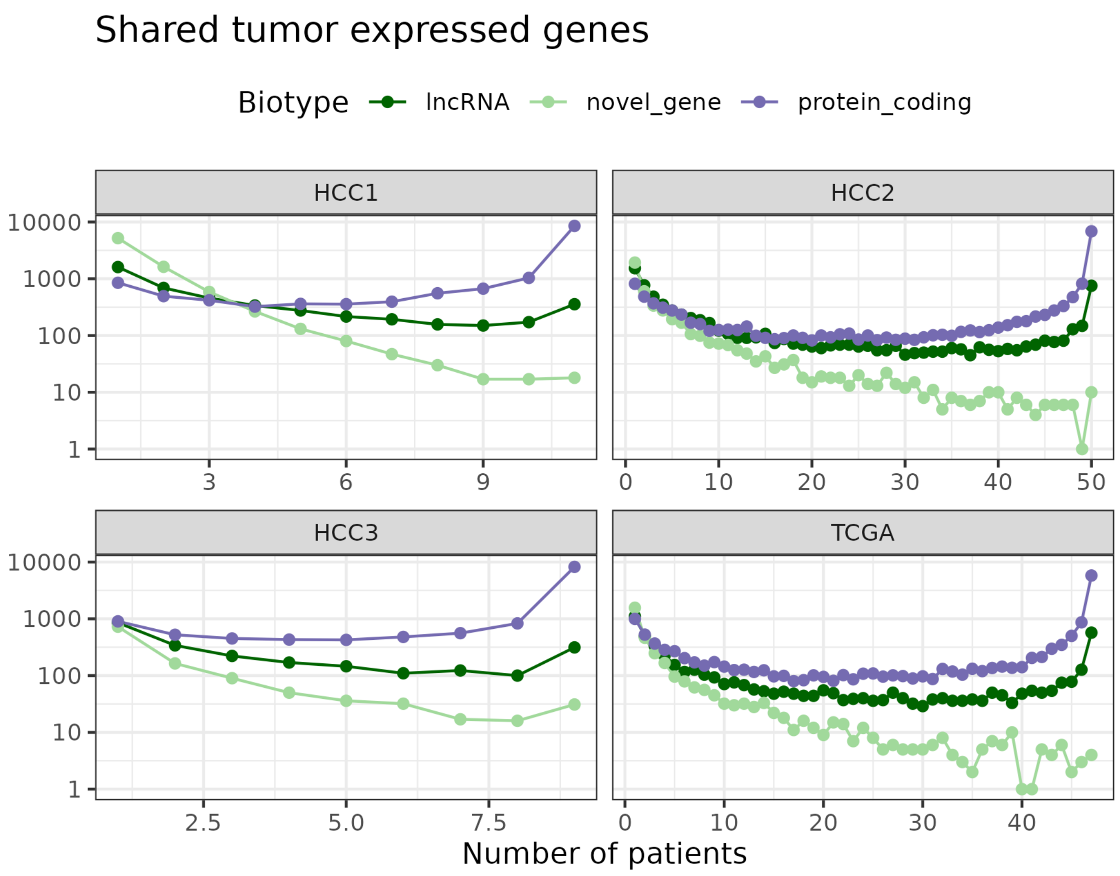

**Figure S6. A RibORF score of 0.5 separates coding sequences and negative controls.** The graph shows the relationship between the number of reads (readNum) and the RibORF score (corrected.pred.pvalue) for different types of transcripts as defined by RibORF. Ribo-Seq data was for dataset HCC4. We selected a cut-off of 0.5 (red line), which divided the main distribution of canonical coding sequences (positive) and that of internal ORFs (negative). The maximum score shown is 1. Both ATG and other near-cognate codons (ACG, CTG, GTG, TTG) were considered as possible start sites. No significant differences in average RibORF score were detected for ORFs initiated at ATG and for ORFs initiated at alternative codons.

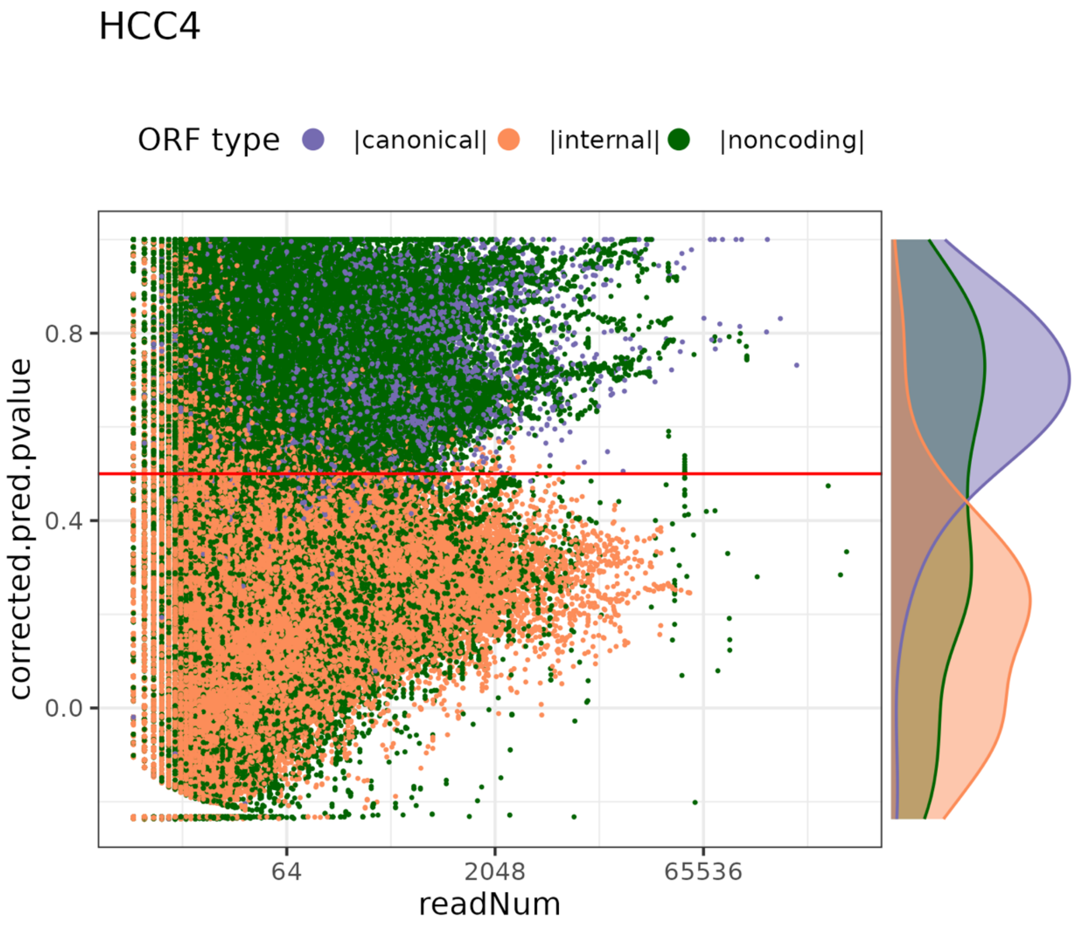

**Figure S7. Translated ncORFs tend to be longer than non-translated ones.** Translated ncORFs refers to those with a RibORF score > 0.5 using data from HCC4. Wilcoxon signed-rank test p-value < 2.2e-16.

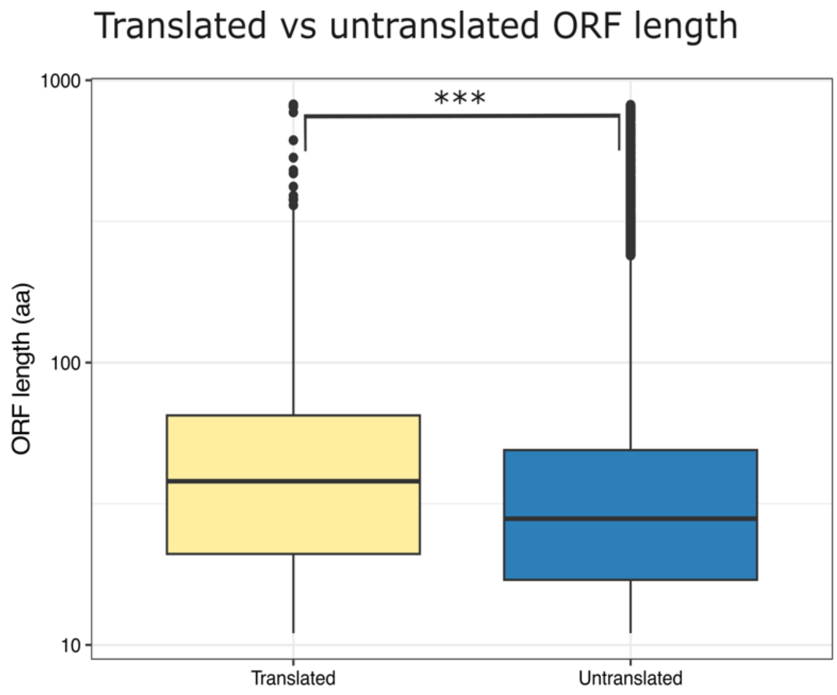

**Figure S8. Number of tumor-specific transcripts per patient and dataset.** LncRNAs and novel transcripts are more prevalent than protein-coding genes.

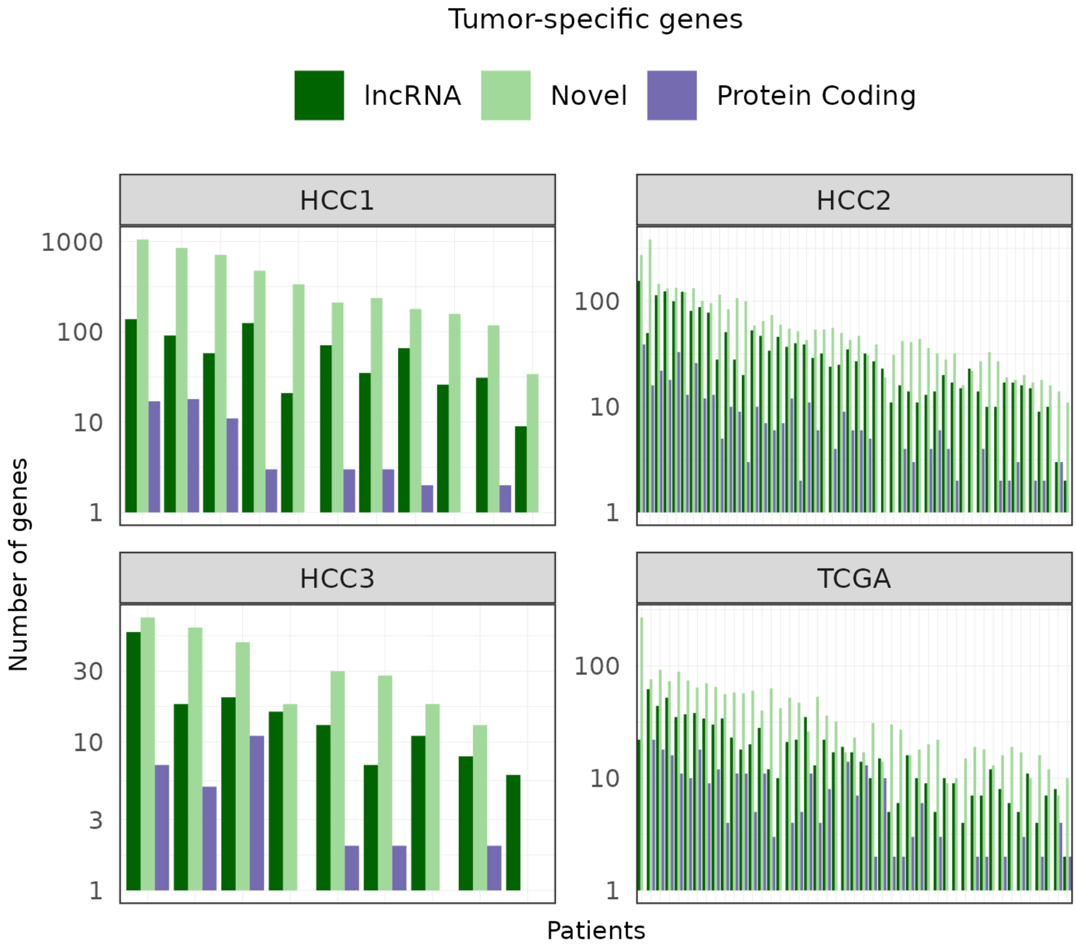

**Figure S9. Comparison between the number of transcripts of different types for tumor-specific and normal-specific transcriptomes.** The tumor-specific transcriptome is computed for each patient by identifying transcripts expressed in a significant manner in the tumor sample but not in the normal sample. The normal-specific transcriptome is computed in the same way but recovering only transcripts that are expressed in the normal sample and not in the tumor sample. Purple: protein-coding genes; dark green: lncRNAs; light green: novel transcripts. The excess of lncRNAs and novel transcripts over protein-coding genes is only consistently observed in the tumor-specific transcriptome.

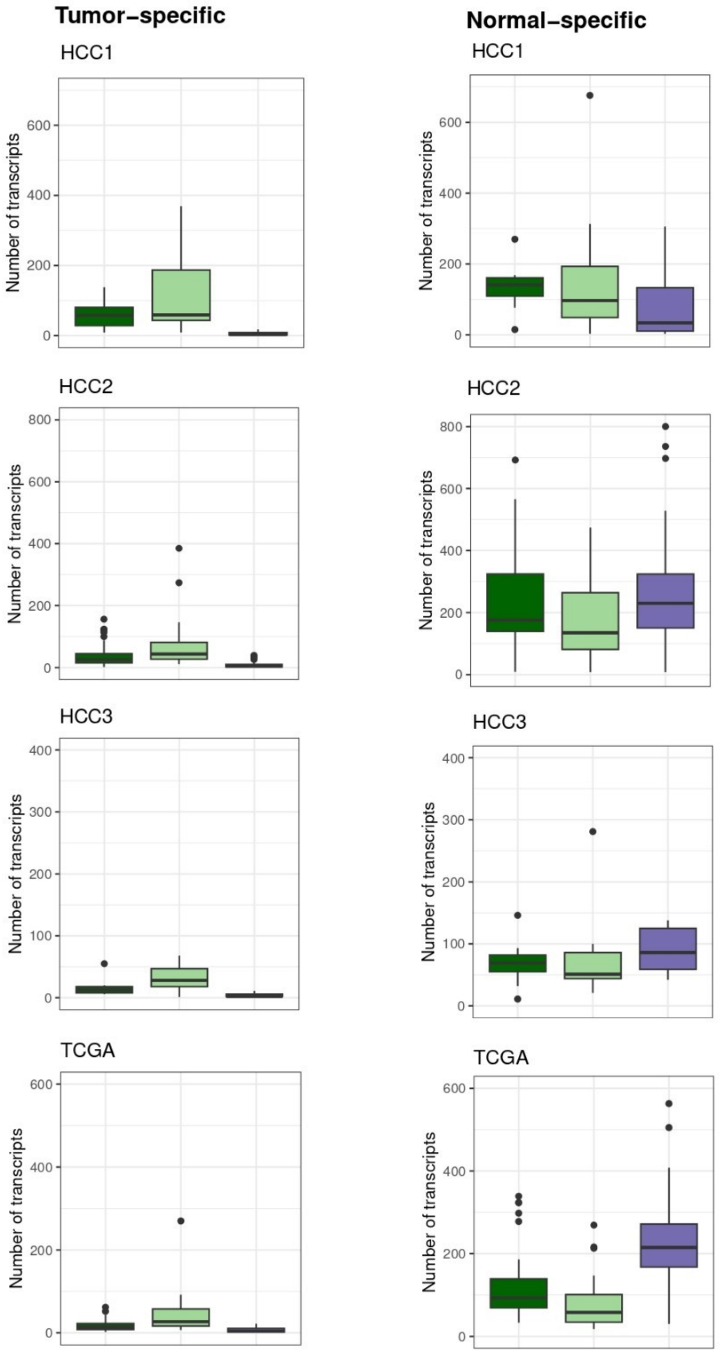

**Figure S10. Proportion of tumor-specific transcripts among all the expressed ones.** The proportion of tumor-specific lncRNAS and novel transcripts is higher than the proportion of tumor-specific protein-coding genes in all datasets.

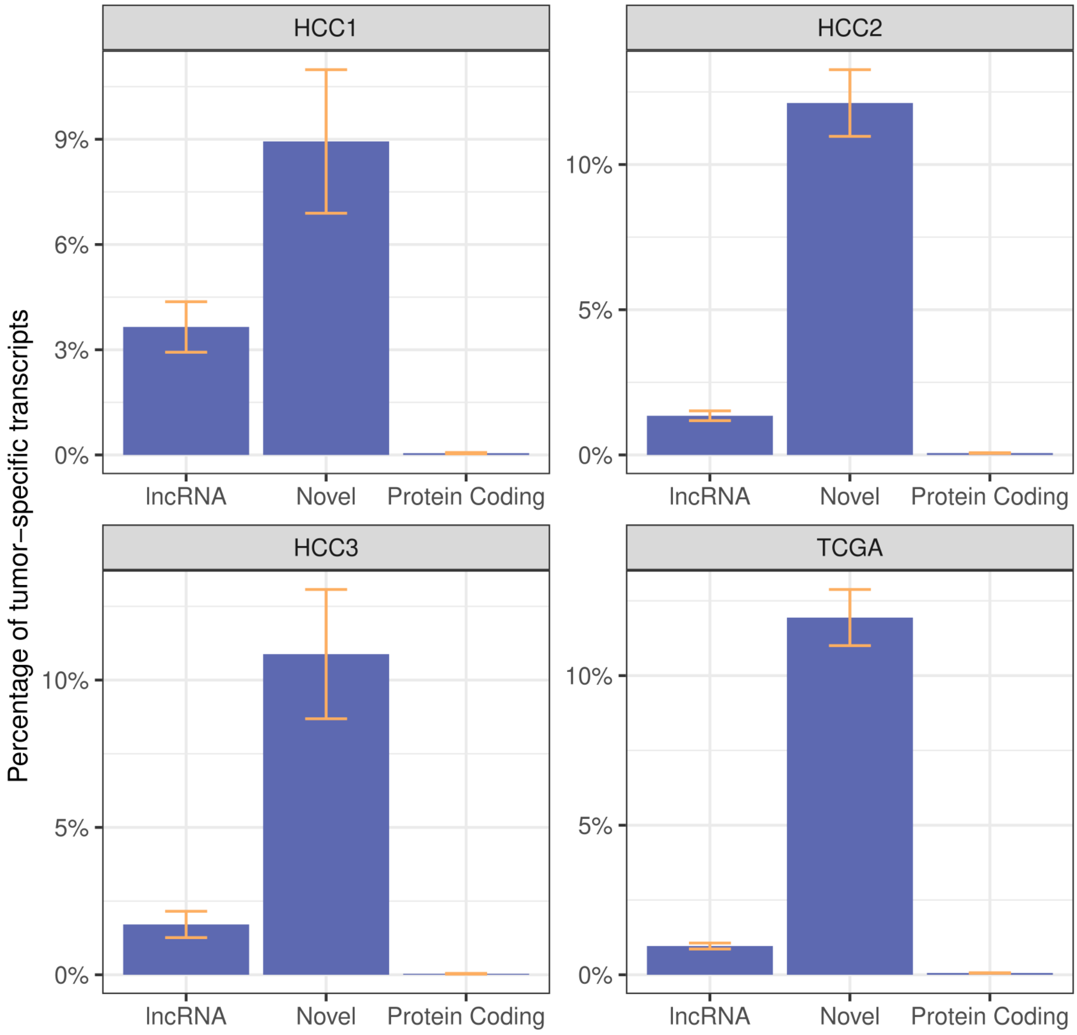

**Figure S11. Expression in testis of different types of tumor-specific transcripts.** Number of transcripts expressed in testis using different expression cut-offs. Protein-coding transcripts are more frequently found in testis, denoting expression in germinal cells, than novel transcritps, with lncRNAs showing intermediate levels. Data for HCC3 is shown in the main manuscript file.

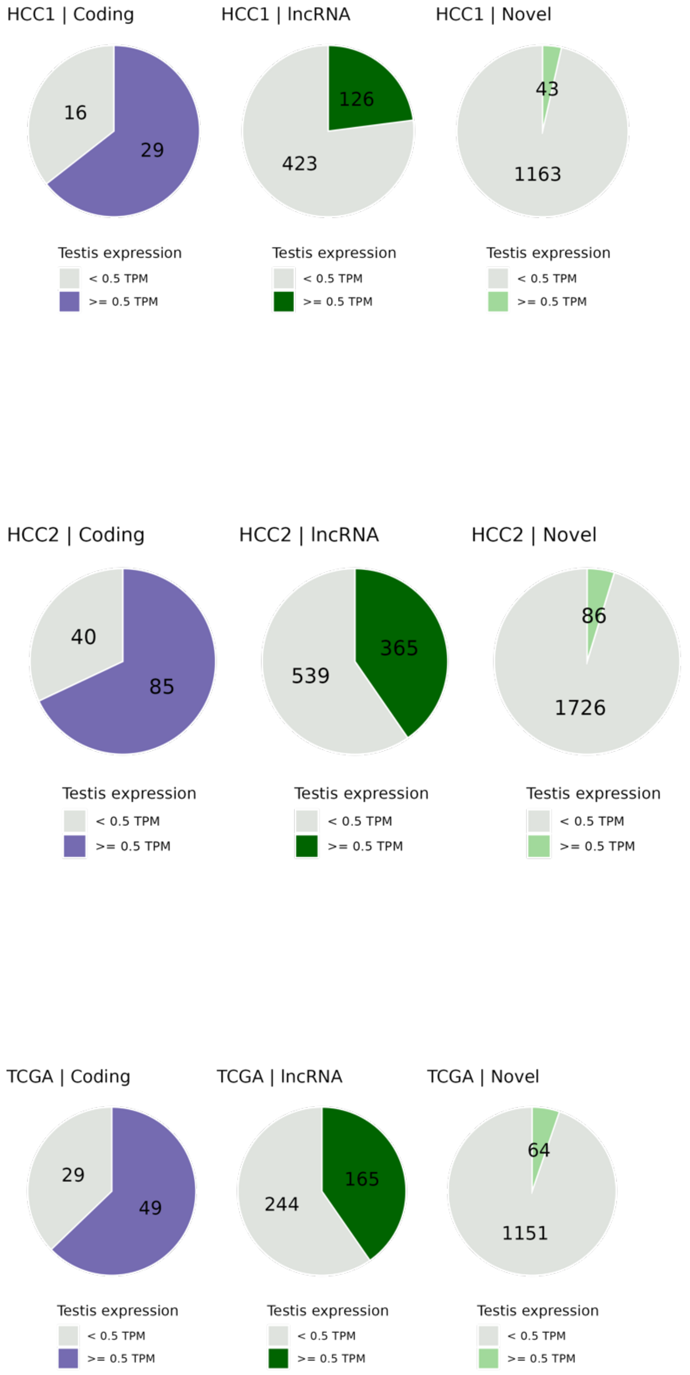

**Figure S12. Validation of tumor-specific gene expression in an external hepatocellular carcinoma cohort.** Gene expression data from an independent HCC cohort (LICA) comprising 161 patient tumor samples was used to measure the patient representativity of the transcripts described in the 117 HCC cohort. Of the genes found in > 10% of the patients in the latter cohort, two were not found in the LICA tables (LINC00221 and BX276092.9). Of the remaining genes, 68% were significantly expressed in at least 10% of the patients in the LICA cohort.

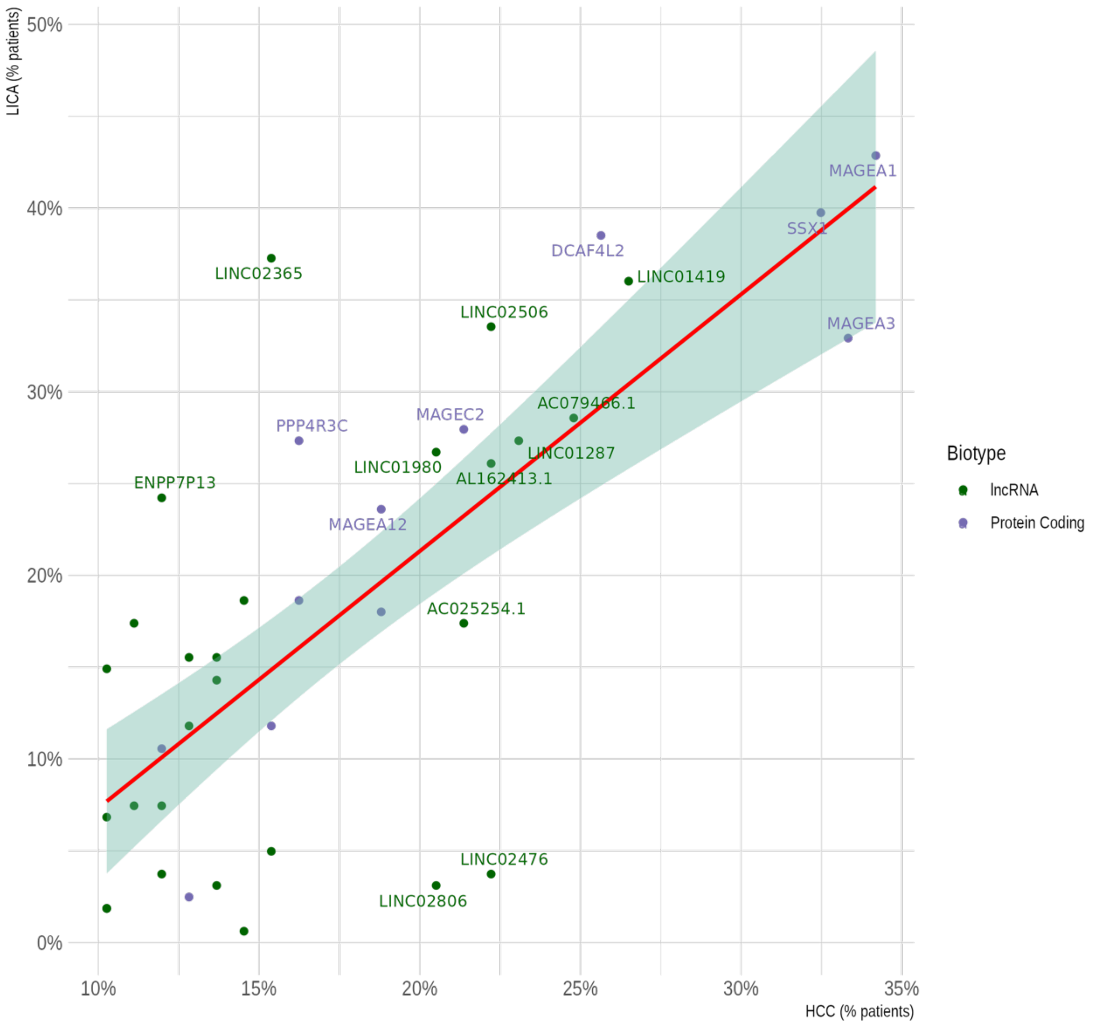

**Figure S13. Correlation of the gene expression values of tumor-specific transcripts in different patients.** Data is for genes expressed in > 10% of the patients in the complete 117 patient cohort. Higher values represent a higher tendency for the two transcripts to be expressed in the same set of patients. Protein-coding genes (cancer/testis antigens) are shown in purple, whereas lncRNAs are shown in black.

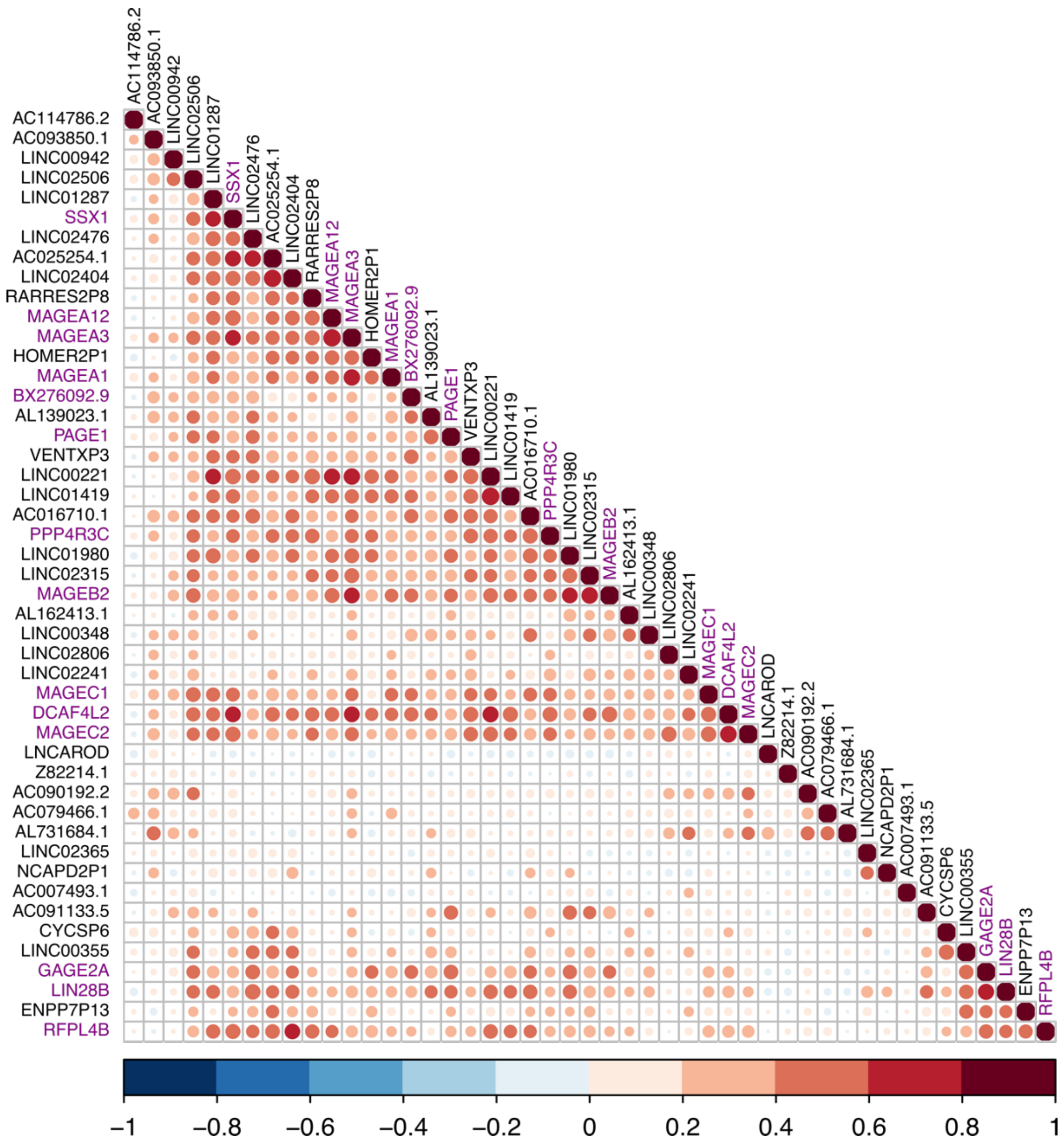
